# Chloride oscillation in pacemaker neurons regulates circadian rhythms through a chloride-sensing WNK kinase signaling cascade

**DOI:** 10.1101/2021.07.16.452737

**Authors:** Jeffrey N. Schellinger, Qifei Sun, John M. Pleinis, Sung-Wan An, Jianrui Hu, Gaëlle Mercenne, Iris Titos, Chou-Long Huang, Adrian Rothenfluh, Aylin R. Rodan

## Abstract

Central pacemaker neurons regulate circadian rhythms and undergo diurnal variation in electrical activity in mammals and flies. In mammals, circadian variation in the intracellular chloride concentration of pacemaker neurons has been proposed to influence the response to GABAergic neurotransmission through GABA_A_ receptor chloride channels. However, results have been contradictory, and a recent study demonstrated circadian variation in pacemaker neuron chloride without an effect on GABA response. Therefore, whether and how intracellular chloride regulates circadian rhythms remains controversial. Here, we demonstrate a signaling role for intracellular chloride in the *Drosophila* ventral lateral (LN_v_) pacemaker neurons. In control flies, intracellular chloride increases in LN_v_ neurons over the course of the morning. Chloride transport through the sodium-potassium-2-chloride (NKCC) and potassium-chloride (KCC) cotransporters is a major determinant of intracellular chloride concentrations. *Drosophila melanogaster* with loss-of-function mutations in the NKCC encoded by *Ncc69* have abnormally low intracellular chloride six hours after lights on, and a lengthened circadian period. Loss of *kcc*, which is expected to increase intracellular chloride, suppresses the long-period phenotype of *Ncc69* mutant flies. Activation of a chloride-inhibited kinase cascade, consisting of the WNK (With No Lysine (K)) kinase and its downstream substrate, Fray, is necessary and sufficient to prolong period length. Fray activation of an inwardly rectifying potassium channel, Irk1, is also required for the long-period phenotype. These results indicate that the NKCC-dependent rise in intracellular chloride in *Drosophila* LN_v_ pacemaker neurons restrains WNK-Fray signaling and overactivation of an inwardly rectifying potassium channel to maintain normal circadian period length.

## INTRODUCTION

Central pacemaker neurons are master regulators of organismal circadian rhythms, which govern physiological processes throughout the body in anticipation of daily changes in light and temperature. In mammals, the central pacemaker neurons reside in the suprachiasmatic nucleus (SCN) of the hypothalamus. Neuronal activity and electrophysiological parameters, such as the resting membrane potential, change in a circadian manner in these neurons (Allen et al., 2017; Harvey et al., 2020). Most SCN neurons are GABA (*γ*-amino butyric acid)-ergic (Ono et al., 2018). One mechanism for GABAergic neurotransmission is through the GABA_A_ receptor, a ligand-gated chloride channel. Because intracellular chloride in neurons is typically low, GABA is usually hyperpolarizing, due to chloride influx (Kaila et al., 2014). However, Wagner *et al*. showed that GABA had excitatory effects on SCN neurons during the day, and inhibitory effects at night, which they proposed was due to higher intracellular chloride concentrations during daytime compared to night (Wagner et al., 1997). Since then, several papers have inferred oscillating intracellular chloride concentrations, as estimated by the GABA reversal potential (Alamilla et al., 2014; Shimura et al., 2002). However, SCN neuron responses to GABA in the day vs. night have differed across studies (Alamilla et al., 2014; Choi et al., 2008; Gribkoff et al., 1999; Irwin and Allen, 2009; Jeu and Pennartz, 2002; Shimura et al., 2002). More recently, using a transgenic chloride sensor to directly measure intracellular chloride concentrations in two specific SCN neuronal subpopulations, Klett and Allen demonstrated increased intracellular chloride during the day, and decreased intracellular chloride at night, as initially proposed by Wagner *et al*. However, GABA application consistently resulted in chloride influx, suggesting hyperpolarizing (inhibitory) effects of GABA in both day and night (Klett and Allen, 2017). Thus, the physiological role of intracellular chloride oscillations in pacemaker neurons remains unclear.

The SLC12 cation-chloride cotransporters are major determinants of intracellular chloride (Kaila et al., 2014). These electroneutral cotransporters couple transmembrane chloride transport to sodium and/or potassium, with chloride influx through the sodium-potassium-2-chloride (NKCC) cotransporters and efflux through the potassium-chloride (KCC) cotransporters. Prior studies have demonstrated SCN neuron expression of these transporters, and physiological roles in determining intracellular chloride and the GABA reversal potential in the SCN (Alamilla et al., 2014; Belenky et al., 2010; Choi et al., 2008; Irwin and Allen, 2009; Klett and Allen, 2017; Shimura et al., 2002).

Here, we use *Drosophila melanogaster*, in which mechanisms regulating circadian rhythms have been extensively studied (Rosbash, 2021; Young, 2018), to understand the role of intracellular chloride in pacemaker neurons. We demonstrate that intracellular chloride increases over the course of the morning in the ventral lateral (LN_v_) pacemaker neurons, in an NKCC-dependent manner. We find that intracellular chloride has a signaling role in these neurons, restraining excess activity of the chloride-sensitive WNK (With No Lysine (K)) kinase, an upstream activator of an inwardly-rectifying potassium channel. Dysregulation of this pathway prolongs the circadian period of diurnal locomotor activity patterns.

## RESULTS

### Intracellular chloride increases in sLN_v_ pacemaker neurons over the course of the morning in an NKCC-dependent manner

The *Drosophila* brain contains ∼150 clock neurons, with the small LN_v_ (sLN_v_) neurons controlling circadian locomotor rhythms by driving the morning peak of activity (King and Sehgal, 2020; Nitabach and Taghert, 2008). sLN_v_ neurons receive input from other neurons, but existing evidence indicates that this does not occur through neurotransmitter-gated chloride channels. In the fly, these include glutamatergic, glycinergic, histaminergic and GABA_A_ chloride channels (Frenkel et al., 2017; Knipple and Soderlund, 2010). Glutamate and GABA signal to adult sLN_v_ neurons through metabotropic receptors, whereas knockdown of the *Rdl* GABA_A_ receptor in LN_v_ neurons has minimal effects on period length (Chung et al., 2009; Collins et al., 2014; Dahdal et al., 2010). While the sLN_v_ neurons themselves are glycinergic (Frenkel et al., 2017), glycinergic signaling to sLN_v_ neurons has not been described, and histaminergic chloride channels are not expressed in sLN_v_ neurons (Hong et al., 2006). Therefore, we reasoned that studying the sLN_v_ neurons would allow us to examine the role of intracellular chloride independent of its effects on the driving force for chloride flux through neurotransmitter-gated chloride channels.

We used a transgenic sensor, ClopHensor, to measure intracellular pH and chloride concentrations in the sLN_v_ neurons *ex vivo* every 4 hours after lights on (ZT0) (Arosio et al., 2010; Mukhtarov et al., 2013; Pleinis et al., 2021; Sun et al., 2018). Calibration curves are shown in Supplemental Figure 1. Intracellular pH, which influences chloride measurement (Arosio et al., 2010), varied by less than 0.06 across time (Supplemental Table 1). Intracellular chloride rose during daytime, fell, and then rose again during nighttime (Supplemental Figure 2). However, brains were exposed to light during dissection and imaging, which could confound nighttime measurements. Still, as in the SCN, intracellular chloride varies in the sLN_v_ pacemaker neurons over the day/night cycle.

In most cell types, activity of the Na^+^/K^+^-ATPase results in low intracellular sodium and high intracellular potassium. NKCCs, therefore, transport chloride along the sodium gradient into the cell, while KCCs typically transport chloride along the potassium gradient out of the cell (Kaila et al., 2014). We therefore measured sLN_v_ intracellular chloride concentrations in flies with strong loss-of-function mutations in *Ncc69*, which encodes an NKCC (Leiserson et al., 2010; Rodan et al., 2012; Sun et al., 2009). pH was similar between control and *Ncc69* mutants (Supplemental Table 1). Intracellular chloride variation in the sLN_v_ neurons was also similar in *Ncc69^r2^* mutants compared to control, with the notable exception that intracellular chloride did not increase from ZT2 to ZT6 (Supplemental Figure 2). We therefore repeated pH and chloride measurements at ZT2 and ZT6 in the same brains in a paired fashion. For control flies, intracellular pH was 7.17±0.01 at ZT2 and 7.14±0.01 at ZT6. For *Ncc69* mutant flies, intracellular pH was 7.16±0.01 at ZT2 and 7.14±0.01 at ZT6. There was no effect of genotype on pH (p=0.9185, two-way repeated measures ANOVA). In controls, intracellular chloride rose from ZT2 to ZT6, whereas chloride concentrations remained constant in *Ncc69* mutants, resulting in lower chloride concentrations in *Ncc69^r2^* mutants compared to controls at ZT6 (Figure 1A). These results indicate that intracellular chloride increases in sLN_v_ neurons over the course of the morning in an NKCC-dependent manner.

**Figure 1.**
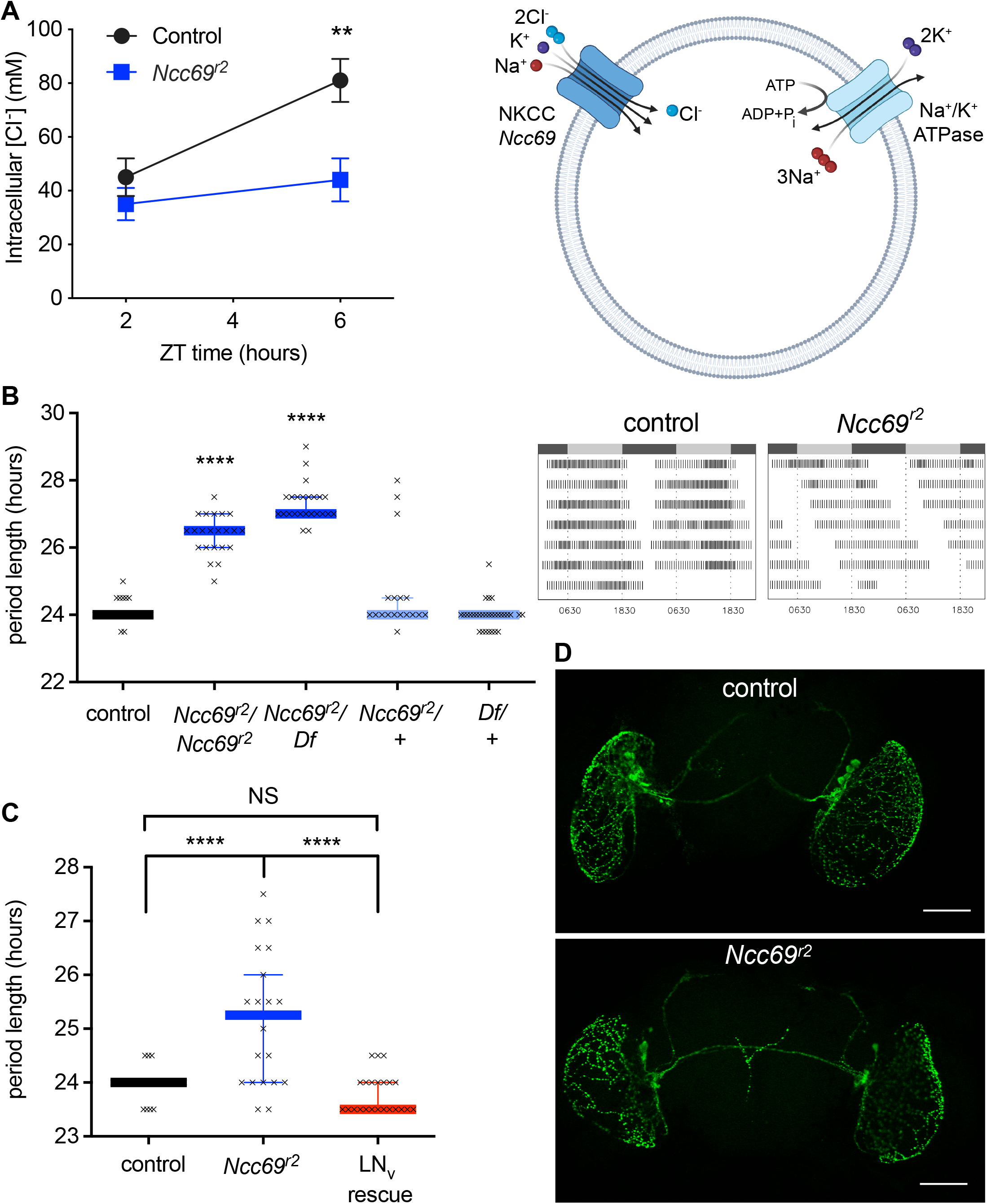
The *Ncc69* NKCC is required in *Drosophila* LN_v_ pacemaker neurons for the morning increase in intracellular chloride and for normal circadian period length. A) Intracellular chloride was measured in sLN_v_ pacemaker neurons using the transgenic sensor, ClopHensor, at 2 and 6 hours after lights-on (ZT2, ZT6). Mean ± SEM shown. Significant effects of time (p=0.0003), genotype (p=0.0125) and interaction (p=0.0166), two-way repeated measures ANOVA. **, p=0.0013 for intracellular chloride in control vs. the *Ncc69^r2^* mutant, Sidak’s multiple comparison test. B) *Ncc69* mutant flies have a prolonged circadian period. In this and subsequent figures, flies were tested for 24-hour locomotor activity in constant darkness for 7 days. Median and 95^th^ percentile confidence intervals with individual data points are shown in this and subsequent figures. In some cases individual data points and/or error bars are obscured by the median line. Complete genotypes and numbers of flies analyzed in this and subsequent figures are listed in Supplemental Table 2. ****, p<0.0001 compared to control, Kruskal-Wallis test with Dunn’s multiple comparisons. Right, representative actograms showing average activity across 7 days of subjective day (grey bars) and night (dark bars) in constant darkness. C) Expression of wild-type *Ncc69* in the LN_v_ pacemaker neurons, under the control of *pdf*-GAL4, rescues the long-period phenotype of *Ncc69* mutant flies. ****, p<0.0001, one-way ANOVA with Tukey’s multiple comparisons test. NS, not significant. D) LN_v_ neuron morphology is intact in *Ncc69* mutants. PDF neuropeptide-expressing pacemaker neurons were visualized using *α*-PDF antibodies. Immunostaining in *Ncc69* mutants and controls were performed in parallel. Scale bar, 100 μm.

### The Ncc69 NKCC is required in LN_v_ pacemaker neurons for normal circadian period

To test the functional consequences of the loss of NKCC activity on circadian rhythms, locomotor activity of individual flies was analyzed in constant darkness using *Drosophila* activity monitors. *Ncc69^r2^* mutant flies had a prolonged period length that was rescued by LN_v_ neuron-specific expression of wild-type *Ncc69* (Figure 1B, C). Although the *pdf-* GAL4 driver used for sLN_v_ neuron expression is also expressed in the large LN_v_ (lLN_v_) neurons (Renn et al., 1999), the latter have been implicated in sleep and arousal and therefore likely do not contribute to the period length phenotype (Chung et al., 2009; Nitabach and Taghert, 2008; Parisky et al., 2008).

As with *Ncc69^r2^* homozygous mutants, prolonged period was also observed in flies in which the *Ncc69^r2^* allele was in trans to a deficiency deleting *Ncc69*, whereas no phenotype was observed in heterozygous mutants (Figure 1B). LN_v_ morphology was intact in *Ncc69^r2^* mutant flies (Figure 1D), as judged by immunostaining with an antibody to the LN_v_-expressed PDF peptide (Park et al., 2000; Renn et al., 1999). We also observed decreased power (weaker rhythmicity) in *Ncc69^r2^* homozygous mutants, but not in *Ncc69^r2^/Df*, and the decreased rhythmicity was not rescued by *Ncc69* expression in the LN_v_ neurons (Supplemental Table 3). Therefore, the reduced rhythm strength may be due to mutation in a different gene, or *Ncc69* activity outside the LN_v_ neurons, and was not pursued further. Roles for *Ncc69* have been demonstrated in glia, including decreased rhythmicity with glial *Ncc69* knockdown (Leiserson et al., 2010; Ng et al., 2016; Rusan et al., 2014; Stenesen et al., 2019). However, expression of wild-type *Ncc69* in glia, using either one or two copies of *gli*-GAL4, did not rescue the long-period phenotype, nor the decreased rhythmicity, of *Ncc69* mutants (Supplemental Table 3). Thus, *Ncc69* is specifically required in the LN_v_ neurons for the maintenance of normal circadian period.

### Intracellular chloride dysregulation results in WNK-Fray kinase activation in the LN_v_ pacemaker neurons

We reasoned that if low intracellular chloride in *Ncc69* mutants is driving the long-period phenotype, decreasing KCC activity could counteract this effect. Period lengths of flies with a heterozygous mutation in *kcc* alone, or with LN_v_ knockdown of *kcc*, were slightly longer than controls (*w/*Y; *kcc^DHS1^/*+, *τ*=25 [24.5-25] hours, median [95% confidence interval], n=15 vs. *w*/Y Berlin, *τ*=24.5 [23.5-26] hours, n=16; *w*/Y; *pdf-*GAL4/*pdf-*GAL4 UAS-kcc^RNAi^, *τ*=24.5 [24-24.5] hours, n=9 vs. *w*/Y; *pdf*-GAL4, *τ*=23.5 [23.5-23.5], n=21). However, heterozygous mutation in *kcc* suppressed the long-period phenotype of *Ncc69* mutants (Figure 2A). This effect is autonomous to the pacemaker neurons, because LN_v_-specific knockdown of *kcc* also suppressed the long-period phenotype of *Ncc69* mutants (Figure 2B). This further supports the idea that dysregulated intracellular chloride is driving the long-period phenotype of *Ncc69* mutants.

**Figure 2.**
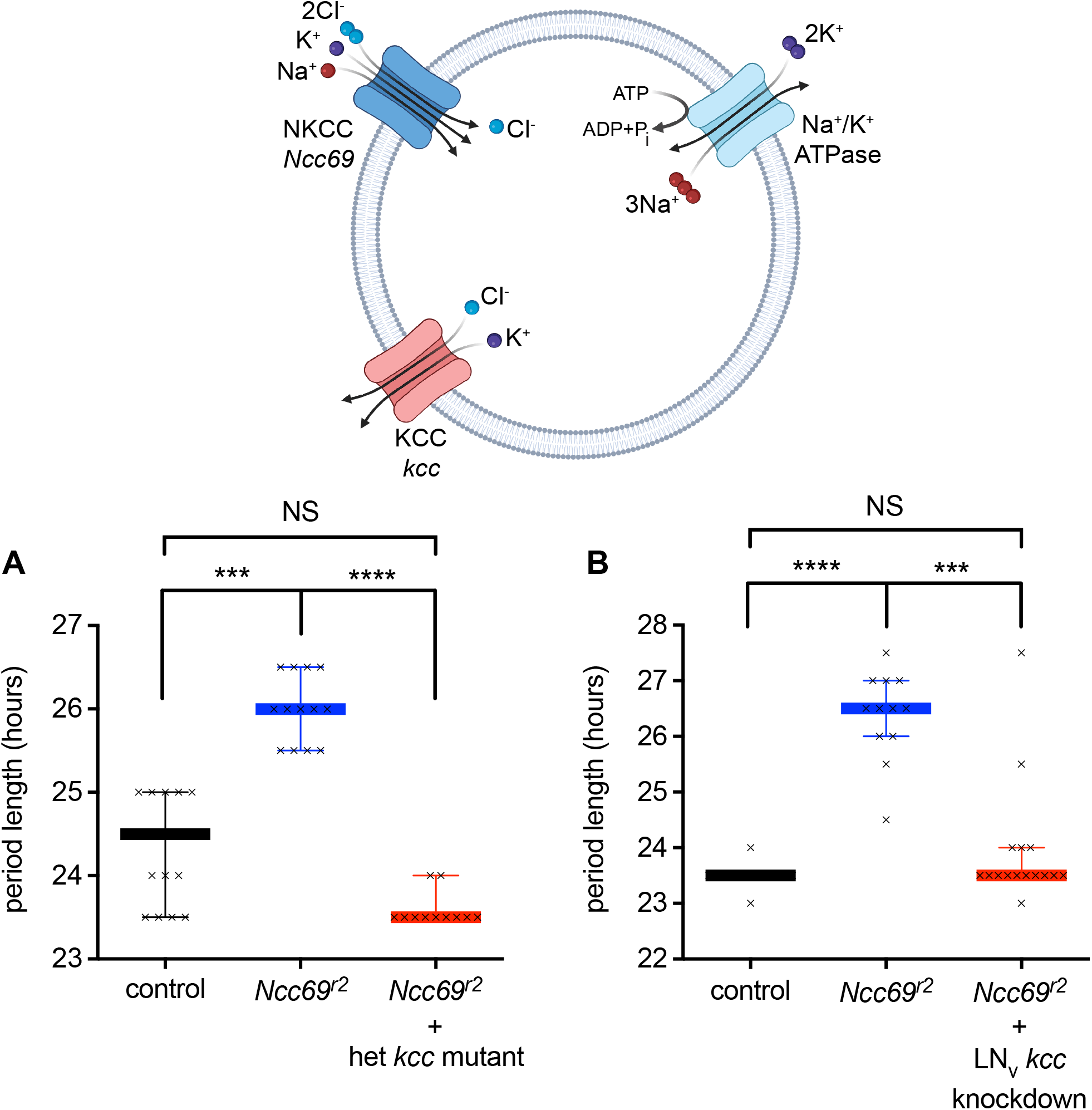
Loss of *kcc* suppresses the long period phenotype of *Ncc69* mutants. A) Global heterozygous loss of *kcc* suppresses the long-period phenotype of *Ncc69* mutants. ***, p=0.0004; ****, p<0.0001, Kruskal-Wallis with Dunn’s multiple comparisons test. B) Knockdown of *kcc* in LN_v_ pacemaker neurons also suppresses the long-period phenotype of *Ncc69* mutant flies. ****, p<0.0001; ***p=0.0001, Kruskal-Wallis test with Dunn’s multiple comparisons.

The *Drosophila* NKCC encoded by *Ncc69* is activated by the WNK-Fray kinase cascade, homologs of the WNK-SPAK/OSR1 (Ste20-related proline alanine rich kinase/oxidative stress response) kinases that activate mammalian NKCCs (Alessi et al., 2014; Rodan, 2018). In addition, this kinase cascade is regulated by chloride, through direct inhibitory effects of chloride ion on mammalian and *Drosophila* WNKs (Piala et al., 2014; Sun et al., 2018; Terker et al., 2016). We first tested the hypothesis that loss of *WNK* or *Fray* in the LN_v_ neurons would phenocopy the *Ncc69* loss-of-function phenotype, i.e. lengthening of the circadian period, due to the loss of positive regulation by the kinases. We knocked down *WNK* in the LN_v_ neurons, using two different UAS-WNK^RNAi^ lines, multiple copies of the *pdf-*GAL4 driver and UAS-WNK^RNAi^, and enhancing the RNAi effect with overexpression of *dicer* (Dietzl et al., 2007). Although we observed a slight increase in period length, this ranged only between 9 and 36 minutes longer than the period length in control flies (Supplemental Table 4). Similarly, *Fray* knockdown, using two different RNAi lines, multiple copies of GAL4/UAS and *dicer* overexpression, had no effect on circadian period length (Supplemental Table 4). Thus, *WNK* and *Fray* do not appear to be acting upstream of *Ncc69* in circadian period regulation.

We next considered the hypothesis that the prolonged circadian period of *Ncc69* mutants is due to the decreased late-morning intracellular chloride in the LN_v_ neurons and excess WNK-Fray activity. Consistent with this hypothesis, LN_v_ neuron knockdown of either *WNK* (Figure 3A) or *Fray* (Figure 3B) suppressed the long-period phenotype of *Ncc69* mutants. Conversely, LN_v_ neuron overexpression of the Cl^-^-insensitive *WNK^L421F^* mutant (Piala et al., 2014; Sun et al., 2018) resulted in circadian period lengthening, phenocopying the *Ncc69* mutant phenotype (Figure 3C). In contrast, there was no effect of overexpressing wild-type *Drosophila WNK*, emphasizing the importance of chloride regulation of WNK in circadian period regulation (Figure 3C). In mammals, WNK3 is expressed in the SCN pacemaker neurons (Belenky et al., 2010). Overexpression of chloride-insensitive human *WNK3^L295F^* in the LN_v_ neurons also resulted in circadian period lengthening, indicating phylogenetically conserved effects of this signaling pathway (Figure 3D). As with *Drosophila WNK*, overexpression of wild-type human *WNK3* resulted in a period length similar to control flies (*w/Y; pdf-*GAL4/UAS-HsWNK3, *τ*=24.5 [24 – 24.5] hours, n=15 vs. *w/Y; pdf-*GAL4/+, *τ*=24 [24 – 24] hours, n=16). These results indicate that loss of WNK chloride inhibition is sufficient to prolong period length.

**Figure 3.**
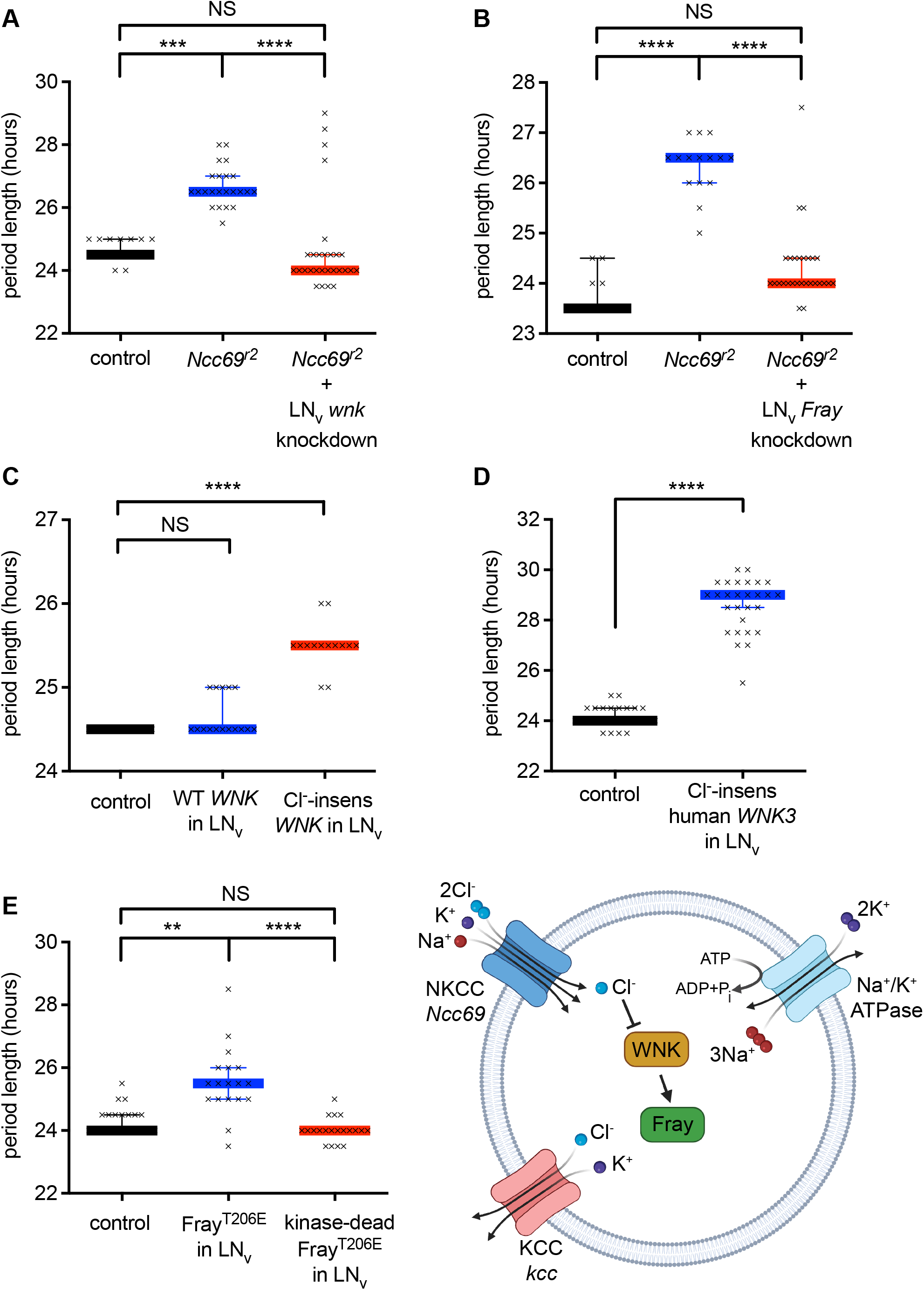
Period lengthening in *Ncc69* mutants is due to low chloride activation of the WNK-Fray kinase cascade. A) Knockdown of the chloride-inhibited *WNK* kinase in LN_v_ pacemaker neurons suppresses the long-period phenotype of *Ncc69* mutants. ***, p=0.0007; ****, p<0.0001, Kruskal-Wallis test with Dunn’s multiple comparisons. B) Knockdown of *Fray*, the kinase downstream of *WNK*, in the LN_v_ neurons suppresses the long-period phenotype of *Ncc69* mutants. ****, p<0.0001, Kruskal-Wallis test with Dunn’s multiple comparisons. C) LN_v_ pacemaker neuron overexpression of Cl^-^-insensitive, but not wild-type, *Drosophila WNK* results in lengthening of circadian period. ****, p<0.0001, one-way ANOVA with Dunnett’s multiple comparisons to control. D) Overexpression of Cl^-^-insensitive human *WNK3* also increases period length. ****, p<0.0001, two-tailed Mann-Whitney test. E) Overexpression of activated Fray^T206E^ increases period length in a kinase-dependent manner; the D185A mutation abolishes kinase activity. **, p=0.0011; ****, p<0.0001, Kruskal-Wallis test with Dunn’s multiple comparisons test.

WNK kinases activate SPAK/OSR1 kinases by phosphorylating the SPAK/OSR1 T-loop threonine (Vitari Biochem J 2005; Anselmo PNAS 2006). We have shown that mutation of the T-loop threonine in Fray, T206, to a phosphomimicking glutamate, results in WNK-independent constitutive activity of Fray *in vitro*, and rescues the ion transport defect in renal tubules in which *Drosophila WNK* is knocked down (Wu et al., 2014). The Fray D185A mutation abolishes kinase activity (Wu et al., 2014). LN_v_ neuron overexpression of *Fray^T206E^* increased period length in a kinase activity-dependent manner, mirroring the phenotype of loss of *Ncc69* or expression of chloride-insensitive WNK (Figure 3E). Period length of the UAS controls was similar to the *w/Y; pdf-*GAL4/+ control (*w/Y;* UAS-Fray^T206E^, *τ*=24.5 [24 – 24.5], n=19; *w/Y;* UAS-Fray^D185A,T206E^, *τ*=24 [23.5 – 24], n=21; *w/Y; pdf-*GAL4/+, *τ*=24 [24-24.5], n=21). Thus, loss of *Ncc69* results in the failure of intracellular chloride to rise during the morning hours, and, as a consequence, WNK and Fray become overactive, initiating a signaling cascade that prolongs circadian period.

### Fray activates the inwardly rectifying potassium channel Irk1 to prolong circadian period

Since the Ncc69 NKCC acts upstream, but not downstream, of the WNK-Fray pathway in the regulation of circadian period length, we were interested in identifying other possible Fray targets required for circadian period prolongation. We queried the *Drosophila* proteome for proteins with putative Fray RFXV/I binding motifs (Delpire and Gagnon, 2007), and identified 127 proteins using an optimized *Drosophila* motif (see Methods) (Supplemental Table 5). We pursued the inwardly rectifying potassium channel, Irk1 (also known as Ir), as a possible target, for three reasons. First, *Irk1* knockdown in the LN_v_ neurons affects circadian period (Ruben et al., 2012). Second, *Irk1* transcript is highly enriched in LN_v_ pacemaker neurons, cycles in a circadian fashion, and also exhibits circadian rhythms in translation (Huang et al., 2013; Kula-Eversole et al., 2010). Third, OSR1 regulates mammalian inwardly rectifying potassium channels containing RFXV-related motifs (i.e., RXFXV) (Taylor et al., 2018).

To examine whether Fray regulates Irk1 channel activity, we performed whole-cell patch clamp recordings of S2-R+ cultured *Drosophila* cells transfected with Irk1, with or without Fray^T206E^ co-expression from a multi-cistronic plasmid. Irk1 activity was determined from barium-sensitive currents (Supplemental Figure 3). Fray^T206E^ expression increased Irk1 channel activity. Mutation of the putative Irk1 Fray-binding RFXV motif to RFXA (Irk1^V306A^) decreased channel activity relative to wild-type Irk1 in the absence of co-transfected Fray^T206E^. Fray is endogenously expressed in S2-R+ cells (Cherbas et al., 2011), so this could represent decreased stimulation by endogenous Fray, although we cannot rule out direct effects of the mutation on the channel. The Irk1 RFXV mutation also blunted stimulation by Fray^T206E^, with no significant difference in Irk1 current density with or without Fray^T206E^ co-expression at a holding potential of -150 mV (Figure 4A, B). Together, these results indicate a stimulatory effect of Fray^T206E^ on wild-type, but not RFXV mutant Irk1.

**Figure 4.**
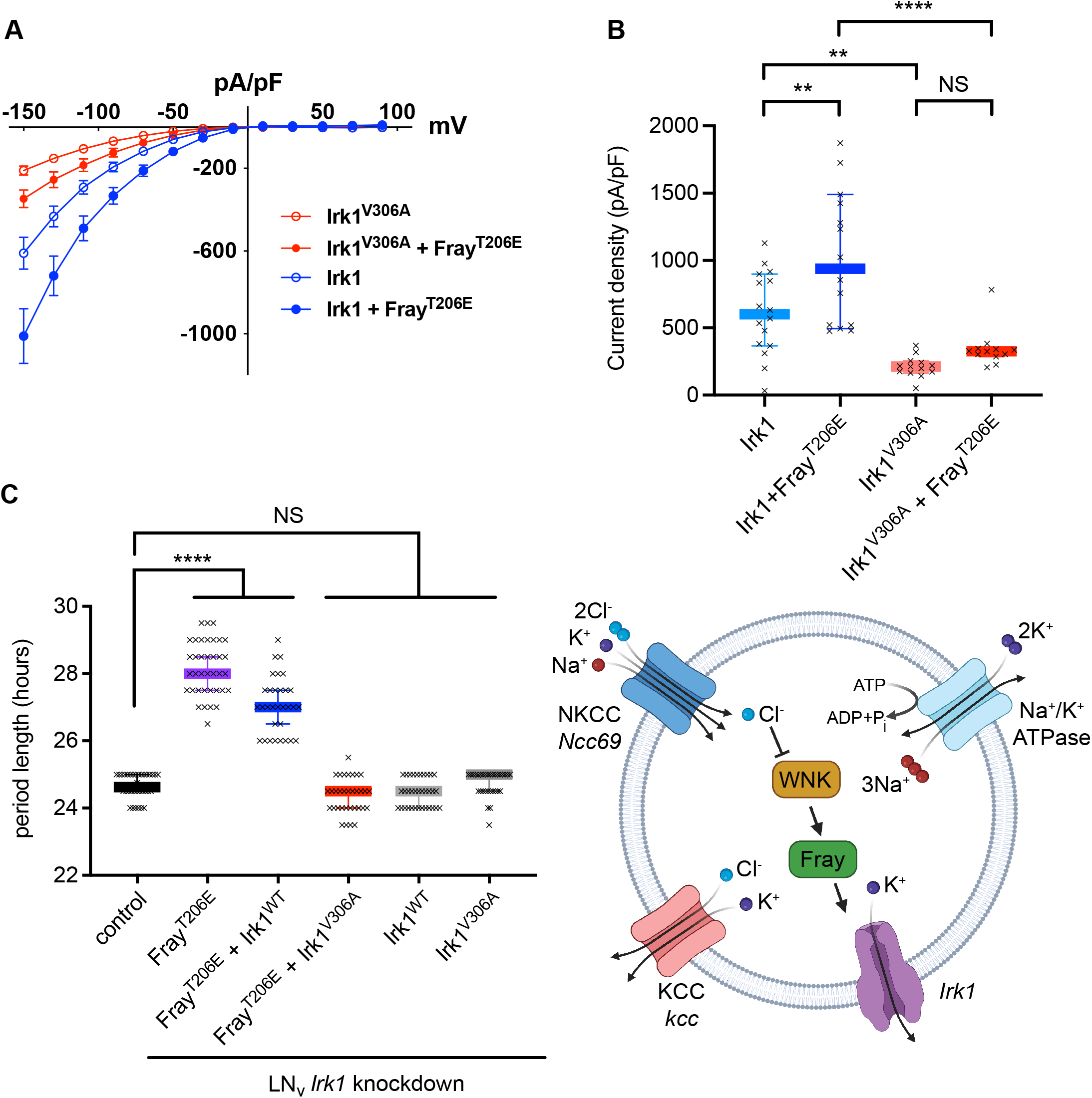
Activated Fray stimulates the inwardly rectifying potassium channel Irk1 to prolong circadian period. Fray^T206E^ expression increases activity of wild-type Irk1, but not Irk1^V306A^, carrying a mutation in the predicted Fray-binding RFXV motif. A) Current-voltage curves from S2-R+ *Drosophila* cultured cells transfected with wild-type Irk1 or Irk1^V306A^, with or without co-expression of constitutively active Fray^T206E^. B) Current density at -150 mV. **, p<0.01, ****, p<0.0001, two-way ANOVA with Tukey’s multiple comparisons test. n=12-16 cells analyzed/genotype. C) Irk1^V306A^ suppresses the long-period phenotype of Fray^T206E^. *Irk1* was knocked down in the LN_v_ pacemaker neurons using RNAi, and replaced with *Irk1^WT^* or *Irk1^V306A^* transgenes with mutations in the RNAi target sites. ****, p<0.0001, Kruskal-Wallis test with Dunn’s multiple comparisons test. A *Drosophila* NKCC regulates circadian rhythms via inhibition of the chloride-sensing WNK kinase and inwardly rectifying potassium channels Jeffrey N. Schellinger, Qifei Sun, John M. Pleinis, Sung-wan An, Jianrui Hu, Gaëlle Mercenne, Iris Titos, Chou-Long Huang, Adrian Rothenfluh, Aylin R. Rodan

We next assessed the *in vivo* role of Fray stimulation of Irk1. We designed Irk1^WT^ and Irk1^V306A^ transgenes in which the sequence targeted by a previously-validated short hairpin RNAi to Irk1 (Wu et al., 2015) was mutated, to allow replacement of endogenous pacemaker neuron Irk1 with either the wild-type or V306A mutant. Knocking down *Irk1* in the pacemaker neurons and replacing it with either *Irk1^WT^* or *Irk1^V306A^* had no effect on period length (Figure 4C). This is consistent with the observation that loss of *Fray* in the pacemaker neurons has no phenotype (Supplemental Table 4). As previously observed, expressing *Fray^T206E^* in the pacemaker neurons increased period length. Flies in which endogenous Irk1 was replaced with Irk1^WT^ in the presence of Fray^T206E^ also had a long-period phenotype. However, replacement of Irk1 with Irk1^V306A^ suppressed the long-period phenotype (Figure 4C). This suggests that Fray stimulation of Irk1 is required for the long-period phenotype observed with loss of *Ncc69* and activation of the WNK-Fray kinase cascade.

## DISCUSSION

Intracellular chloride oscillates in the central pacemaker neurons of the mammalian SCN (Alamilla et al., 2014; Klett and Allen, 2017; Shimura et al., 2002; Wagner et al., 1997), but the functional significance of this oscillation has remained unclear. Here, we demonstrate an NKCC-dependent increase in intracellular chloride in *Drosophila* LN_v_ pacemaker neurons over the course of the morning, which constrains activity of the chloride-sensitive WNK kinase, its downstream substrate, Fray, and an inwardly rectifying potassium channel, Irk1, to maintain normal circadian periodicity.

We show that loss of the Ncc69 NKCC in the LN_v_ pacemaker neurons has two consequences: the failure of intracellular chloride to increase over the course of the morning, and lengthening of the circadian period. Our observations are consistent with studies in the SCN implicating NKCC in the determination of intracellular chloride in mammalian pacemaker neurons (Alamilla et al., 2014; Belenky et al., 2010; Choi et al., 2008; Irwin and Allen, 2009; Klett and Allen, 2017; Shimura et al., 2002), and connects intracellular chloride to a behavioral circadian phenotype. Another SLC12 transporter has been linked to *Drosophila* behavioral rhythmicity in constant light conditions and the electrophysiological response to GABA in lLN_v_ neurons, which have GABA_A_ receptor chloride channels (Buhl et al., 2016; Chung et al., 2009; Parisky et al., 2008). This transporter was named *NKCC* and was proposed to regulate intracellular chloride. However, its transport activity has not been characterized, and it aligns to a distinct clade of SLC12 cotransporters whose sequences predict differences in ion selectivity compared to the NKCCs (Piermarini et al., 2017). Consistent with this idea, an *Aedes aegypti* homolog has a transport profile distinct from the NKCCs, with an electrogenic, chloride-independent lithium/sodium conductance (Kalsi et al., 2019). Thus, the functions of the *Ncc69*-encoded NKCC and the *NKCC-*encoded transporter are likely distinct.

Diurnal variations in intracellular chloride have been proposed to influence the effect of GABAergic neurotransmission on clock neurons in mammals (Alamilla et al., 2014; Choi et al., 2008; Gribkoff et al., 1999; Irwin and Allen, 2009; Jeu and Pennartz, 2002; Shimura et al., 2002; Wagner et al., 1997), but a recent study questioned this idea (Klett and Allen, 2017). Here, we demonstrate a signaling role for intracellular chloride in the LN_v_ neurons, in which the rise in intracellular chloride inhibits WNK-Fray signaling. As chloride-sensitive kinases, WNKs are perfectly poised to interpret changes in intracellular chloride and initiate downstream signal transduction cascades, and the concept of chloride signaling has gained increasing traction in recent years (Lüscher et al., 2020; Piala et al., 2014; Rodan, 2019). This has been studied in the context of transepithelial ion transport in *Drosophila* and mammalian renal epithelia (Chen et al., 2019; Sun et al., 2018; Terker et al., 2015), as well as in the clearance of apoptotic corpses (Perry et al., 2019). Our finding that chloride signals via the WNK-Fray pathway in circadian pacemaker neurons further extends this concept.

We also show that Fray activates the Irk1 inwardly rectifying potassium channel, in a manner dependent on a putative Fray-binding motif in Irk1 that is required for the lengthened circadian period of flies with activated Fray in LN_v_ neurons. Pacemaker neurons in both flies and mammals undergo molecular clock-controlled circadian variation in electrical activity (Allen et al., 2017; Harvey et al., 2020; King and Sehgal, 2020), and altering the excitability of the LN_v_ pacemaker neurons disrupts circadian rhythms (Depetris-Chauvin et al., 2011; Mizrak et al., 2012; Nitabach et al., 2002, 2006; Ruben et al., 2012; Wu et al., 2008). Variation in pacemaker neuron resting membrane potential is thought to be an important determinant of the circadian pattern of electrical activity (Allen et al., 2017; Harvey et al., 2020). Two voltage-gated potassium channels have been implicated in LN_v_ neuron electrical oscillations (Smith et al., 2019), and a sodium leak current and potassium channels contribute to the day/night cycling of resting membrane potential in *Drosophila* dorsal clock neurons (Flourakis et al., 2015). Because inwardly rectifying potassium channels play an important role in determining cellular membrane potential (Hibino et al., 2010), chloride regulation of Irk1 activity could also contribute to the diurnal variation in LN_v_ neuron excitability, theraby influencing circadian period.

In sLN_v_ neurons, resting membrane potential is most depolarized at lights on (ZT0), hyperpolarizes between ZT0 and ZT6, is variable between ZT6 and ZT18, and depolarizes between ZT18 and ZT24 (Cao and Nitabach, 2008). Thus, the increased intracellular chloride we observe in sLN_v_ neurons at ZT6, which is predicted to inhibit Irk1 through inhibition of WNK-Fray signaling, coincides with the period in which there is loss of hyperpolarization in the sLN_v_ neurons. Persistent Irk1 activity, e.g. in *Ncc69* mutants, could maintain sLN_v_ neurons in a hyperpolarized state, delaying subsequent depolarization over the course of the night and prolonging the circadian period. In contrast, the loss of Irk1 throughout the day-night cycle, as seen with *Irk1* knockdown in the PDF-expressing neurons (Ruben et al., 2012), could interfere with sLN_v_ neuron hyperpolarization after lights-on, explaining the long-period phenotype of those mutants.

Our results imply that intracellular chloride may regulate pacemaker neuron excitability in a cell autonomous fashion. This is consistent with the observation that circadian rhythms of pacemaker neuron electrophysiology persist in the absence of fast neurotransmission and in isolated neurons in invertebrates and vertebrates (Jeu et al., 1998; Michel et al., 1993; Welsh et al., 1995). This raises the question of how NKCC activity may be increasing during morning hours in the LN_v_ neurons. Although the WNK-SPAK/OSR1/Fray kinase cascade is a well-understood upstream regulator of NKCCs, we show that in the LN_v_ neurons, NKCC is acting upstream of WNK, and loss of *WNK* and *Fray* in the pacemaker neurons does not recapitulate the long-period phenotype of loss of *Ncc69*. This suggests other regulatory mechanisms.

Could intracellular chloride play a signaling role in SCN pacemaker neurons? NKCC1, KCCs and WNK3 are expressed in the rat SCN (Belenky et al., 2010). The repertoire of ion channels regulating pacemaker neuron excitability is complex and incompletely understood, but large-conductance Ca^2+^-activated potassium channels (also known as BK or maxi-K) have been implicated, and are regulated by mammalian WNKs (Harvey et al., 2020; Liu et al., 2014; Ray et al., 2021; Wang et al., 2013; Webb et al., 2016; Yue et al., 2013; Zhuang et al., 2011). Whether the oscillating intracellular chloride observed in SCN neurons regulates these or other ion channels underlying SCN neuron electrical properties will be of interest.

## METHODS

### *Drosophila* Strains and Fly Husbandry

*Drosophila melanogaster* strains used are shown in Supplemental Table 6 and the genotypes used for each experiment are shown in Supplemental Table 2. *w; pdf-*GAL4, *w*; UAS-WNK^RNAi^, *w*; UAS-Fray^RNAi^, *w*; UAS-Irk1^RNAi^, *w;* UAS-kcc^RNAi^, *w;* UAS-Fray^T206E^, *w;* UAS-WNK^WT^, *w*; UAS-WNK^L421F^, *w;* UAS-WNK3^WT^, *w;* UAS-WNK3^L295F^, *w;* UAS-Fray^D185A,T206E^ and *w;* UAS-dcr-2 were outcrossed for 5 generations to *wBerlin*, which was also used for generating heterozygous controls (e.g., *w; pdf-*GAL4/+). Except for *w;* UAS-Fray^RNAi^ (Vienna), and *w;* UAS-kcc^RNAi^, knockdown of the targeted genes has previously been validated by qRT-PCR, as referenced in Supplemental Table 6. Recombinant chromosomes and combinations of transgenes were generated by standard genetic techniques. Generation of new transgenic strains is described below. Flies were reared on a standard cornmeal-yeast-molasses medium prepared in a central kitchen at UT Southwestern or the University of Utah. Flies were reared at room temperature (22-23 °C) or at 25 °C. Young adult (<2 week old) male flies were used in all assays. Males were used to avoid female egg-laying and the emergence of larvae in *Drosophila* activity monitors.

### Generation of transgenics

The open reading frame encoding Fray^D185A,T206E^ was recombined from pENTR-Fray^D185A,T206E^ (Wu et al., 2014) into the pUASg.attB Gateway-compatible destination vector, obtained from Johannes Bischof and Konrad Basler (Zürich, Switzerland) (Bischof et al., 2007), using LR Clonase II (Thermo Fisher Cat #11791020). After sequence confirmation, midiprep DNA was injected into stock #24481 (*y^1^M{vas-int.Dm}ZH-2A w*\*; *M{3xP3-RFP.attP′}ZH-22A*) by Rainbow Transgenic Flies (Camarillo, CA). Single male transformants were isolated by the presence of ‘mini white’ and confirmation of the UAS-transgene was performed by PCR with sequence-specific primers.

A plasmid (FI16807, stock #1644763) containing the open reading frame of *Irk1* was obtained from the *Drosophila* Genomics Resource Center (Indiana University, Bloomington, IN). The open reading frame was PCR-amplified (Phusion high-fidelity DNA polymerase, New England Biolabs Cat #M0530) using primers Irk1-F and Irk1-R (Supplemental Table 7). After gel extraction (Qiagen QIAquick Cat #28104), the PCR product was cloned into pENTR using the pENTR/D-TOPO cloning kit (Thermo Fisher Cat #K240020) and the sequence confirmed by Sanger sequencing. Next, a mutation was introduced into pENTR-Irk1^WT^ (Supplemental Table 8) to generate a mutation in the putative Fray-binding motif, in which Val 306 in the “RFXV” motif is mutated to Ala. The corresponding “GTG” was mutated to “GCG” using QuikChange II XL (Agilent Cat #200521) and primers Irk1-V306A-F and Irk1-V306A-R to generate pENTR-Irk1^V306A^ (Supplemental Table 8). Next, mutations were introduced into pENTR-Irk1^WT^ and pENTR-Irk1^V306A^ to render the transgenes resistant to knockdown by co-expression of the Irk1 RNAi. Every third nucleotide in the twenty base pairs targeted by the RNAi was mutated in order to introduce five silent mutations (i.e., CTAAAGGAACGCTTC was mutated to CTGAAAGAGCGTTTT), using QuikChange II XL (Agilent Cat #200521) and primers Irk1-RR-F and Irk1-RR-R. The resulting plasmids were called pENTR-Irk1^WT-RR^ and pENTR-Irk1^V306A-RR^ (Supplemental Table 8). Irk1 sequences in all plasmids were confirmed by Sanger sequencing. The open reading frames of Irk1^WT-RR^ and Irk1^V306A-RR^ were then recombined into pUASg.attB using LR Clonase II (Thermo Fisher Cat #11791020) to generate pUASg.attB-Irk1^WT-RR^ and pUASg.attB-Irk1^V306A-RR^ (Supplemental Table 8). Midiprep DNA was injected into stock #24483 (*M{vas-int.Dm}ZH-2A, M{3xP3-RFP.attP}ZH-51D*), transformants isolated, and UAS-transgenes confirmed as above.

### Circadian rhythm analysis

Male flies were collected within 72 hours of eclosion and kept on standard food for 3-5 days in a 12-hour oscillating light/dark incubator. After the entrainment period, individual flies were placed into 5 mm diameter glass cuvettes with standard medium at one end and a tissue plug at the other, allowing the flies free movement throughout the cuvette. Beam breaking by locomotor activity was recorded in 30 minute increments (bins) using Drosophila Activity Monitors (TriKinetics) in constant darkness over a period of 7 days. Activity data were then analyzed using FAASx software (Paris-Saclay Institute of Neuroscience). Cycle p analyses provided period length (tau) and rhythmic strength (power) for individual flies. Settings used were: Minimum period peak power 20, Minimum period peak width 0200. Output was restricted to flies with period lengths of 26 hours ± 10 hours (minimum tau 16, maximum 36). Flies that did not survive the full 7 days (168 hours) or were arrhythmic (power less than 20) were not included in tau analysis.

For the Irk1 experiment, analysis was performed in ClockLab version 6 (Actimetrics, Wilmette, IL), using the chi squared periodogram function. Settings used were: Start Day 1, End Day 9, type chi squared, start 16 hours (minimum tau), end 36 hours (maximum tau), significance 0.01. In order to classify rhythmic strength, cutoffs were determined by assaying *wBerlin* control flies. Flies that did not survive the full 8 days or were arrhythmic (power less than 2500) were not included in tau analysis.

### PDF neuron immunohistochemistry

Brains were dissected from adult flies in PBS (phosphate-buffered saline, in mM: 137 NaCl, 2.7 KCl, 8.1 Na_2_HPO_4_, 1.5 KH_2_PO_4_, pH 7.3-7.4) and fixed for 20-40 minutes in 4% formaldehyde. Brains were then rinsed in PBS, followed by PBT (PBS+0.3% Triton X-100) 4-5 times. Brains were incubated in mouse anti-PDF (PDF C7 from Developmental Studies Hybridoma Bank, Iowa City, IA, (Cyran et al., 2005)), 1:800 in 10% normal goat serum in PBT, overnight at 4°C, then rinsed 4-5 times in PBT and 2-3 times in PBS. Brains were then incubated in 1:800 goat anti-mouse Alexa Fluor 488 secondary antibody (Thermo Fisher Scientific Cat #A-11001) overnight, rinsed in PBT and PBS, and then mounted in 80% glycerol in PBS. Imaging was performed using a Zeiss LSM510 confocal microscope.

### Measurement of intracellular Cl^-^ concentration

#### pH calibration

pcDNA3-ClopHensor (Addgene plasmid #25938) was cut with HindIII and NotI and the resulting product ligated into pIB (Thermo Fisher Cat #V802001) to generate pIB-ClopHensor (Supplemental Table 8). pIB-ClopHensor was transiently transfected into S2-R+ cultured *Drosophila* cells obtained from Helmut Krämer (UT Southwestern) using CellFectin reagent (Thermo Fisher Cat #10362100). 48 hours after transfection, cells were bathed in pH varied solution containing: 38 mM Na gluconate, 100 mM K gluconate, 0.6 mM MgSO_4_, 20 mM HEPES (varied pH), 10 μM tributyltinchloride (Sigma Cat #T50202), 5 μM nigericin (Thermo Fisher Cat #N1495), 5 μM carbonyl cyanide 3-chlorophenylhydrazone (Sigma Cat # C2759) and 5 μM valinomycin (Sigma Cat # V0627). After equilibration for at least 1 hour, cells were imaged using a Zeiss LSM510 confocal microscope, with excitation at 488 nm (green emission) and 458 nm (cyan emission). Individual cells (19-25 cells for each pH) were then outlined and pixel intensity measured using ImageJ without image manipulation. The ratios of green/cyan vs. pH were entered into GraphPad Prism, and a sigmoidal curve interpolated using the function “sigmoidal, 4PL, X is log(concentration).” This provided the values for the following equation, used to calculate intracellular pH (*pH_i_*) in the pacemaker neurons (Mukhtarov et al., 2013):

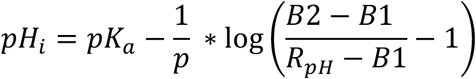

where *R_pH_* is the experimentally derived green/cyan ratio, *pK_a_* = 7.254, *p* = power (Hill slope, 1.668), and *B1* (0.2603) and *B2* (1.915) are the minimum and maximum asymptotic values of *R_pH_*.

#### Cl^-^ calibration

Brains expressing ClopHensor in the pacemaker neurons (*w/Y; pdf-*GAL4 UAS- ClopHensor c202) were dissected from 3-5 day old flies in *Drosophila* saline, consisting of (in mM): NaCl 117.5, KCl 20, CaCl_2_ 2, MgCl_2_ 8.5, NaHCO_3_ 10.2, NaH_2_PO_4_ 4.3, HEPES 15, and glucose 20, pH 7.0. Brains were attached to the bottom of 35 mm glass bottom dishes with 14 mm microwell/#1.5 cover glass (Cellvis) coated with poly-lysine, and the solution exchanged to the chloride calibration solution, consisting of (in mM): 100 mM Na-Cl/gluconate, 50 mM K-Cl/gluconate, 2 mM Ca-Cl/gluconate, 8.5 mM Mg-Cl/gluconate, 20 mM glucose, 15 mM HEPES pH 7.1, 10 μM tributyltinchloride, 5 μM nigericin, 5 μM carbonyl cyanide 3-chlorophenylhydrazone and 5 μM valinomycin. Cl/gluconate anions were adjusted to achieve varying chloride concentrations. After 1 hour equilibration, brains were imaged using a Zeiss LSM510 confocal microscope, with excitation at 543 nm (red emission) and 458 nm (cyan emission). Individual neuron cell bodies (10 per Cl^-^ concentration, from multiple brains) were outlined and pixel intensity measured in ImageJ without image manipulation. The ratios of cyan/red vs Cl^-^ were entered into GraphPad Prism, and a sigmoidal curve interpolated using the function “sigmoidal, 4PL, X is log(concentration).” This provided the values for the following equation, used to calculate intracellular Cl^-^ ([Cl^-^]_i_) (Mukhtarov et al., 2013):

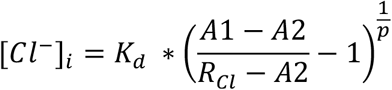

where *R_Cl_* is the experimentally derived cyan/red ratio, *K_d_* = 53.49, *p* = power (set to 1 per methods of (Mukhtarov et al., 2013)), and *A1* (1.538) and *A2* (0.597) are the maximum and minimum asymptotic values of *R_Cl_*.

#### Measurement of R_pH_ and R_Cl_

Male flies were entrained in 12:12 L:D (light:dark) conditions for 4 days. Brains expressing ClopHensor in the pacemaker neurons were removed from the incubator at specific ZT times (time after lights on), after which they were exposed to ambient daytime light, and then dissected in the following solution, adapted from solutions used for two-photon calcium imaging of fly brain neuronal activity (Wang et al., 2003): in mM, NaCl 108, KCl 5, CaCl_2_ 2, MgCl_2_ 8.2, NaHCO_3_ 4, NaH_2_PO_4_ 1, trehalose 5, sucrose 10, HEPES 5, pH 7.5. Dissected brains were attached to the bottom of 35 mm glass bottom dishes with 14 mm microwell/#1.5 cover glass (Cellvis) coated with poly-lysine, and then bathed in the above solution for about 60 minutes prior to imaging using a Zeiss LSM510 confocal microscope, with excitation at 543 nm (red emission), 488 nm (green emission), and 458 nm (cyan emission). Individual sLN_v_ neuronal cell bodies (distinguished from lLN_v_ neurons based on position and size) were outlined in ImageJ and pixel intensity captured for each emission channel. The ratios of green/cyan and cyan/red were used to calculate pH and Cl^-^ as described above. pH and Cl^-^ were measured for each individual neuron. For practical reasons, measurements on brains removed at different ZT times across 24 hours were performed on separate days, but control and *Ncc69* mutant brains were tested in parallel. However, when comparing the ZT2 and ZT6 timepoints, flies were removed from the incubator at ZT2, brains dissected and imaged as above, and then the same brains were re-imaged 4 hours later without further manipulation.

### Proteome-wide search for Fray binding motifs

The *Drosophila* proteome was searched for putative Fray binding motifs, using the methods of Delpire and Gagnon (Delpire and Gagnon, 2007). First, the NCBI protein database was searched for *Drosophila melanogaster* and results were saved to a FASTA text file. Duplicate results and results from organisms other than *D. melanogaster* were eliminated. This list was then searched for two motifs. One motif, [S/G/V]RFx[V/I]xx[I/V/T/S], was derived from Delpire and Gagnon (Delpire and Gagnon, 2007), and is called the “Gagnon motif” in Supplemental Table 5. However, this screen failed to identify Ncc69, which we have previously validated as a Fray target (Wu et al., 2014). It also failed to identify KCC, which is a validated mammalian SPAK/OSR1 target (Heros et al., 2014; Melo et al., 2013; Piechotta et al., 2002; Zhang et al., 2016). We therefore performed sequence alignment of the *Drosophila* homologs of three of the best-known families of mammalian SPAK/OSR-interacting proteins: WNKs, NKCCs/NCC, and KCCs (*Drosophila* WNK, Ncc69, and *Drosophila* KCC, respectively). From this we derived a second motif, referred to as the “*Drosophila* motif” in Supplemental Table 5: [D/E/N/Q/S/T/Y]RFx[V/I]xxxx[D/E/G/P]. A second list of proteins was generated using this motif.

### Patch clamp analysis of Fray^T206E^ effects on Irk1

#### Generation of plasmids for S2 cell expression

To generate plasmids for cellular expression of Irk1^WT^ and Irk1^V306A^, with or without Fray^T206E^ co-expression, the Irk1^WT^ and Irk1^V306A^ open reading frames were amplified from pENTR-Irk1^WT^ or pENTR-Irk1^V306A^ by PCR (Phusion high-fidelity DNA polymerase, New England Biolabs Cat #M0530). Primers pAc5-Irk1-F and pAc5-Irk1-R included additional sequence for subsequent Gibson assembly cloning into the multi-cistronic vector, pAc5 STABLE2 Neo (Addgene Cat #32426, (González et al., 2011)), which uses T2A sequences to generate multiple polypeptides off of a single transcript, and also contains a GFP cassette. The pAc5 plasmid backbone was PCR-amplified using primers pAc5-F and pAc5-R. The products were then assembled using NEBuilder HiFi DNA Assembly (New England Biolabs Cat #E2621) to generate plasmids pAc5-Irk1^WT^ and pAc5-Irk1^V306A^ (Supplemental Table 8). Irk1 inserts were confirmed by Sanger sequencing. Then, the open reading frame encoding Fray^T206E^ was amplified from an existing pENTR-Fray^T206E^ plasmid (Wu et al., 2014), using primers pAc5-Fray-F and pAc5-Fray-R, and the pAc5-Irk1^WT^ or pAc5-Irk1^V306A^ plasmid backbones were amplified using primers pAc5-Irk1-Fray-F and pAc5-Irk1-Fray-R. Using this strategy, the Neo^R^ cassette was replaced by the open reading frame encoding Fray^T206E^, while the GFP cassette was retained to allow for identification of successfully transfected cells. After Gibson assembly, inserts were confirmed by Sanger sequencing. However, the Irk1 ORF contained a stop codon at the end, which would prevent expression of the downstream Fray^T206E^. The stop codon was removed using QuikChange II XL (Agilent Cat #200521) and primers Irk1-TGA-F and Irk1-TGA-R.

#### Patch clamp of Irk1-expressing S2-R+ cells

S2-R+ *Drosophila* cultured cells were obtained from the *Drosophila* Genomics Resource Center (stock #150) and cultured at 25 °C in Schneider’s medium with 10% FBS (Thermo Fisher Cat #10082139). Cells were seeded at a density of 1.7 x 10^6^ cells/ml in 12 well dishes for 24 hours and resuspended in serum-free medium prior to transfection. Cells were transfected with 1 μg plasmid DNA (pAc5-Irk1^WT^, pAc5- Irk1^V306A^, pAc5-Irk1^WT^-Fray^T206E^, or pAc5-Irk1^V306A^-Fray^T206E^) in 100 μL Opti-MEM (Thermo Fisher Cat #31985088) using 2 μL of TransIT-Insect transfection reagent (Mirus Bio Cat #6104). Medium was replaced with serum-containing medium 5 hours after transfection. Irk1 activity was measured in GFP+ cells 48-72 hours after transfection. Cells were harvested by centrifugation and resuspended in fresh medium. They were then plated on cover slips coated with poly-L-lysine (Sigma P8920). Ruptured whole cell recordings were performed in a bath solution containing (in mM) 135 KCl, 1 MgCl_2_, 2 CaCl_2_, 15 glucose, 10 HEPES, 15 sucrose, pH 7.4 with Tris. Patch pipettes were pulled from borosilicate glass capillaries (Sutter Instruments) and heat-polished to give input resistances of 2-3 megaohms. The pipette recording solution contained (in mM) 135 KCl, 1 MgCl_2_, 2 ATP-Mg, 0.1 GTP-Na, 5 EGTA, 10 HEPES, pH 7.2 with Tris. Cells were held at 0 mV and stimulated for 400 ms with step pulses from -150 mV to +90 mV with 20 mV steps. Currents were recorded with an Axopatch 200B patch-clamp amplifier and Pulse software (Molecular Devices, Sunnyvale, CA). 0.5 mM Ba^2+^ inhibited inward currents with 135 mM K^+^, and washing with Ba^2+^-free 135 mM K^+^ recovered the currents. Ba^2+^-sensitive current was therefore analyzed. Data acquisition and analysis were performed using pClamp v.9.2 (Molecular Devices).

### Statistics

Statistical testing was performed using GraphPad Prism, version 9. Data sets were analyzed for normality using the D’Agostino & Pearson normality test. Normally distributed data were compared using t-test or ANOVA and non-normally distributed data were compared using Mann-Whitney or Kruskal-Wallis test. Multiple comparisons testing was performed as indicated in the figure legends. p<0.05 was considered statistically significant. Number of flies or cells examined is indicated in Supplemental Table 2.

## ACKNOWLEDGEMENTS

The authors would like to thank Billy Leiserson, Michael Rosbash and Daria Hekmat-Scafe for fly lines, Johannes Bischof and Konrad Basler for plasmids, and Helmut Krämer for S2-R+ cells. Stocks obtained from the Bloomington *Drosophila* Stock Center (NIH P40OD018537) and the Vienna Drosophila Resource Center were used in this study. Plasmids and S2-R+ cells were obtained from the *Drosophila* Genomics Research Center (Indiana University, Bloomington, IN, supported by NIH grant 2P40OD010949). The PDF C7 antibody, developed by Justin Blau (New York University), was obtained from the Developmental Studies Hybridoma Bank, created by the NICHD of the NIH and maintained at The University of Iowa, Department of Biology, Iowa City, IA 52242. Schematics were drawn using BioRender.

## AUTHOR CONTRIBUTIONS

A.R.R. conceived the study, directed experiments, analyzed data, and wrote the manuscript. J.N.S., Q.S., J.M.P., and S.A. performed experiments. C.L.H. directed the experiments performed by S.A. J.H. and G.M. generated key reagents. I.T. and A.R. provided intellectual input. This work was supported by the National Institutes of Health: DK091316, DK106350 and DK110358 to A.R.R.; DK111542 to C.L.H.; and AA019526, AA026818 and DA049635 to A.R.

## SUPPLEMENTAL INFORMATION

**Supplemental Figure 1, related to Figure 1.**
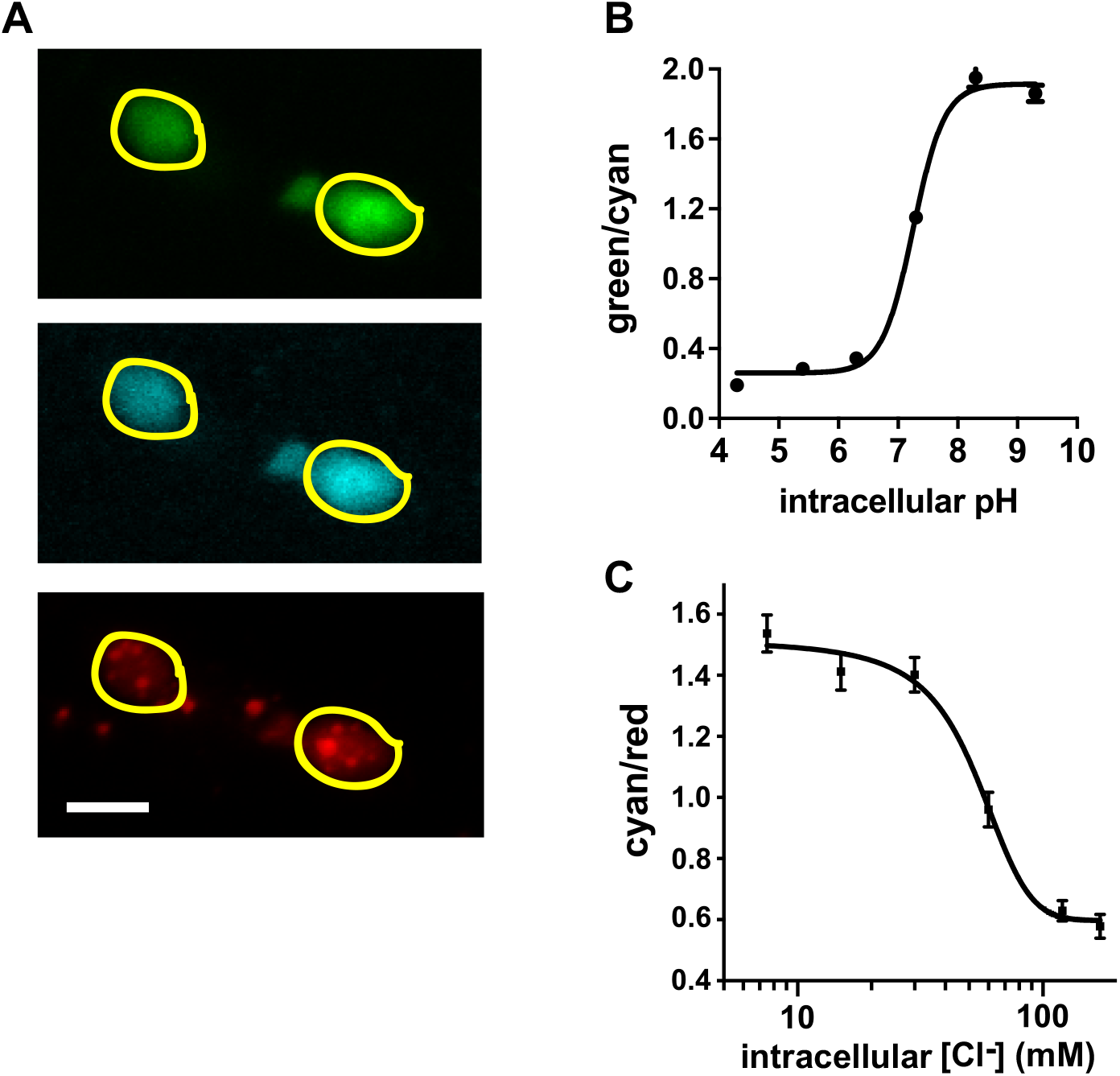
ClopHensor pH and Cl^-^ calibration curves. A) Brains expressing ClopHensor in the LN_v_ pacemaker cells (*w/Y; pdf-*GAL4 UAS-ClopHensor c202) were imaged by confocal microscopy with excitation at 488 nm (green emission), 458 nm (cyan emission) and 543 nm (red emission). A region of interest around each cell to be measured was hand-drawn in ImageJ and used for each of the three channels for measurement of pixel intensity in individual cells (in this case, two cells analyzed). Scale bar = 10 μm. B) pH calibration curve. Cultured S2-R^+^ cells were transiently transfected with ClopHensor. Intra- and extra-cellular pH was equilibrated through the use of tributyltinchloride, nigericin, carbonyl cyanide 3- chlorophenylhydrazone, and valinomycin. pH was varied and the cells were imaged, with pixel intensity measured in ImageJ. The green/cyan ratio at each pH is shown (mean±SEM, note in some cases SEM is smaller than the symbol, n=19-25 cells imaged/pH value), together with the interpolated sigmoidal function. C) Chloride calibration curve. Brains expressing ClopHensor in the PDF-expressing pacemaker neurons (*w/Y; pdf-*GAL4 UAS-ClopHensor c202) were dissected. Intra- and extra-celllular Cl^-^ was equilibrated using the same drug cocktail as above. In preliminary experiments, intracellular pH of the pacemaker neurons was 7.1, therefore the Cl^-^ calibration was performed at pH 7.1. The brains were imaged, with pixel intensity measured in ImageJ. The cyan/red ratio at each Cl^-^ concentration is shown (mean±SEM, n=10 cells imaged/Cl^-^ concentration), together with the interpolated sigmoidal function.

**Supplemental Table 1, related to Figure 1.** pH_i_ of control and *Ncc69* mutants. All values shown are mean±SEM. n=number of neurons tested. ZT0 = lights on. Genotypes: control, *w/Y; pdf-*GAL4 UAS-ClopHensor c202. *Ncc69* mutant, *w/Y; pdf-*GAL4 UAS-ClopHensor c202*; Ncc69^r2^*.

| Circadian<br>time (h) | pH |  |  |  |
| --- | --- | --- | --- | --- |
|  | control | n | *Ncc69*<br>mutant | n |
| ZT2 | 7.15±0.01 | 25 | 7.15±0.01 | 24 |
| ZT6 | 7.17±0.01 | 32 | 7.15±0.01 | 30 |
| ZT10 | 7.19±0.01 | 29 | 7.14±0.01 | 26 |
| ZT14 | 7.19±0.01 | 24 | 7.14±0.01 | 24 |
| ZT18 | 7.21±0.01 | 28 | 7.19±0.01 | 30 |
| ZT22 | 7.21±0.01 | 30 | 7.19±0.01 | 30 |

**Supplemental Figure 2, related to Figure 1.**
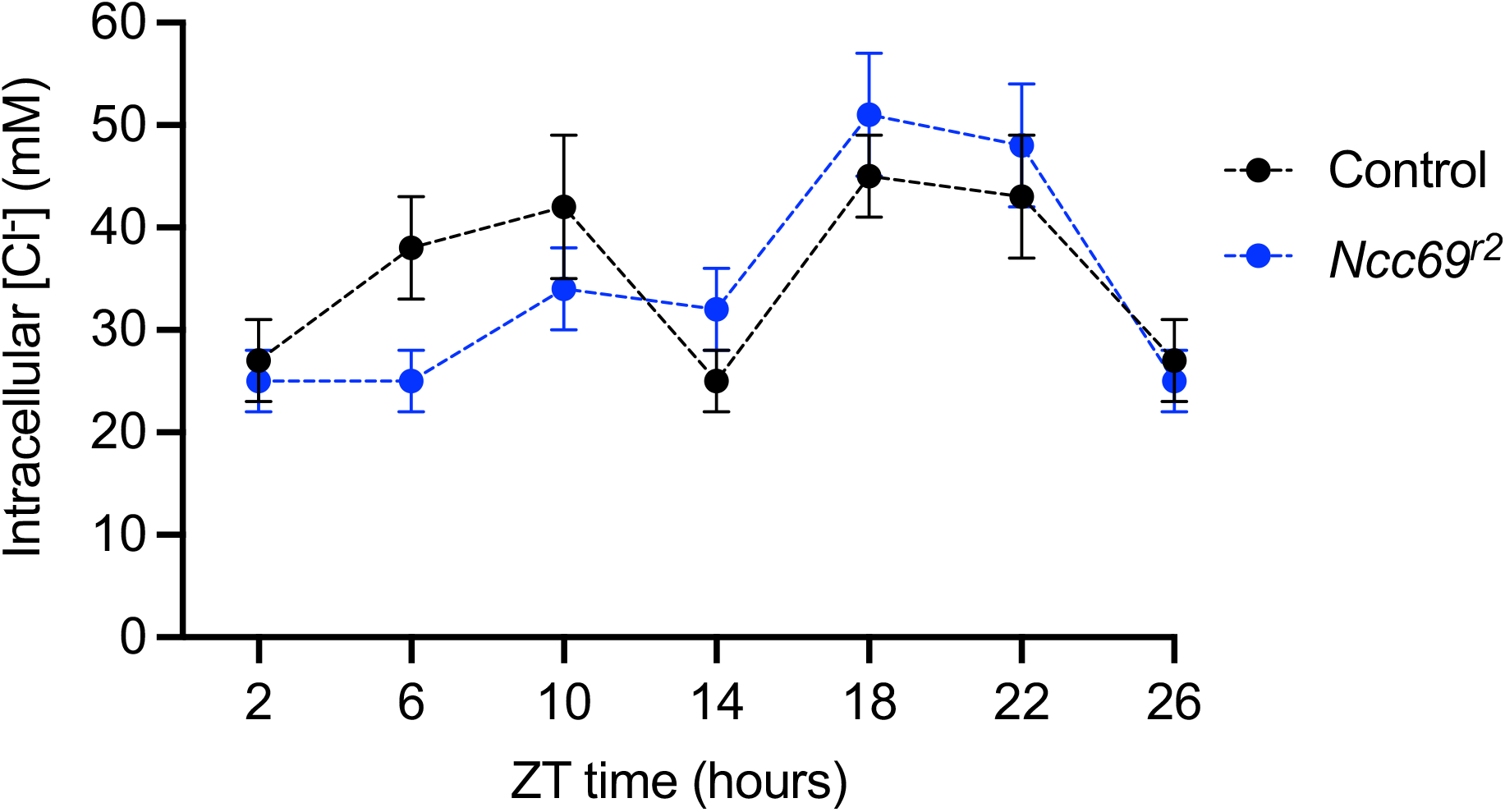
[Cl^-^]_i_ of control and *Ncc69* mutants. Brains expressing ClopHensor in the LN_v_ pacemaker cells (*w/Y; pdf-*GAL4 UAS-ClopHensor c202 and *w/Y; pdf-*GAL4 UAS-ClopHensor c202*; Ncc69^r2^*) were imaged by confocal microscopy as described in Supplemental Figure 1 to obtain the cyan/red ratio, from which intracellular Cl^-^ concentration was calculated using the calibration curve in Supplemental Figure 1. Intracellular pH from the same neurons is shown in Supplemental Table 1, which also includes the number of neurons sampled for each genotype and timepoint. Mean ± SEM shown. ZT2 is double-plotted as ZT26. p<0.0001 for time, p=0.5978 for genotype, two-way ANOVA.

**Supplemental Table 2, related to Figures 1-4.**
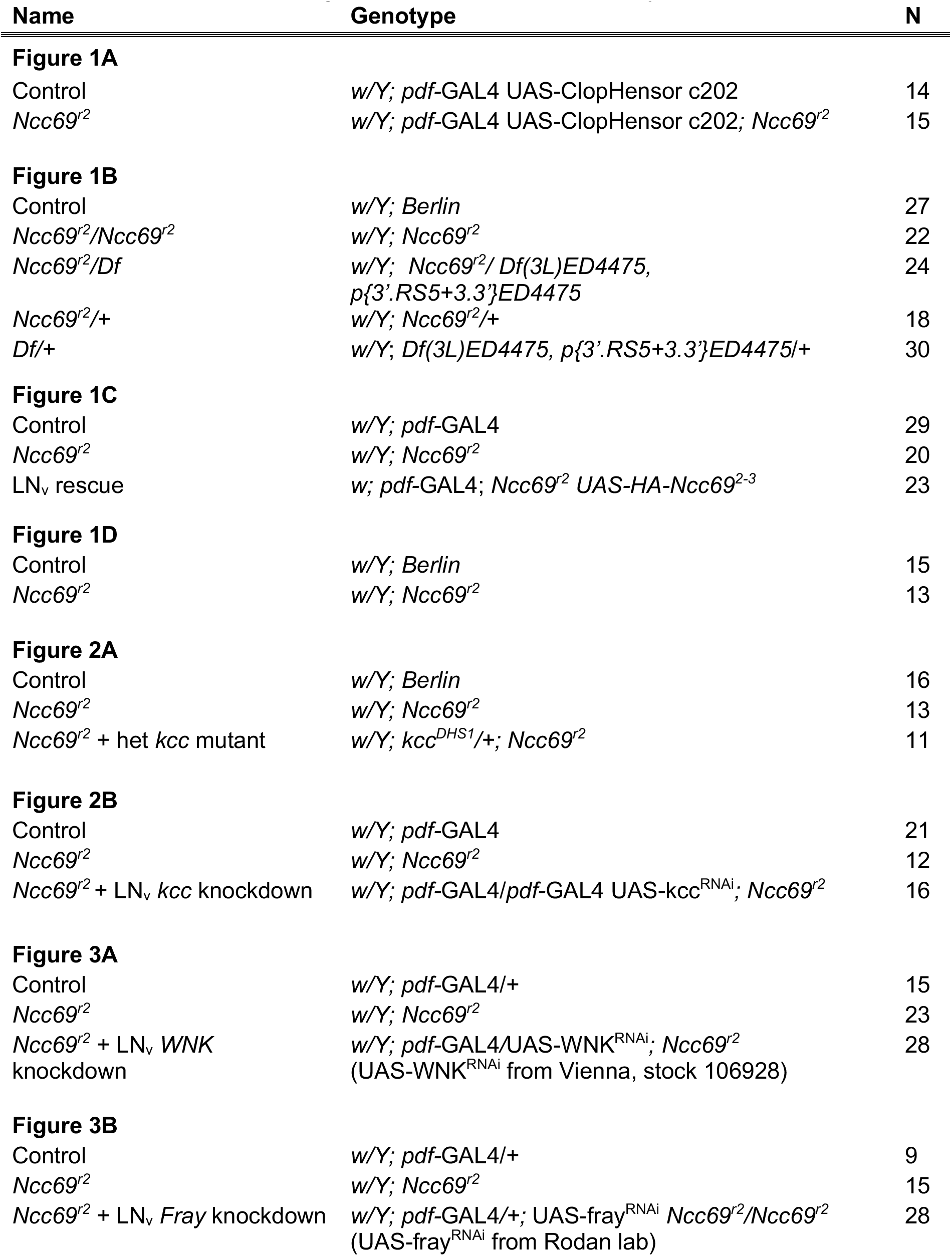
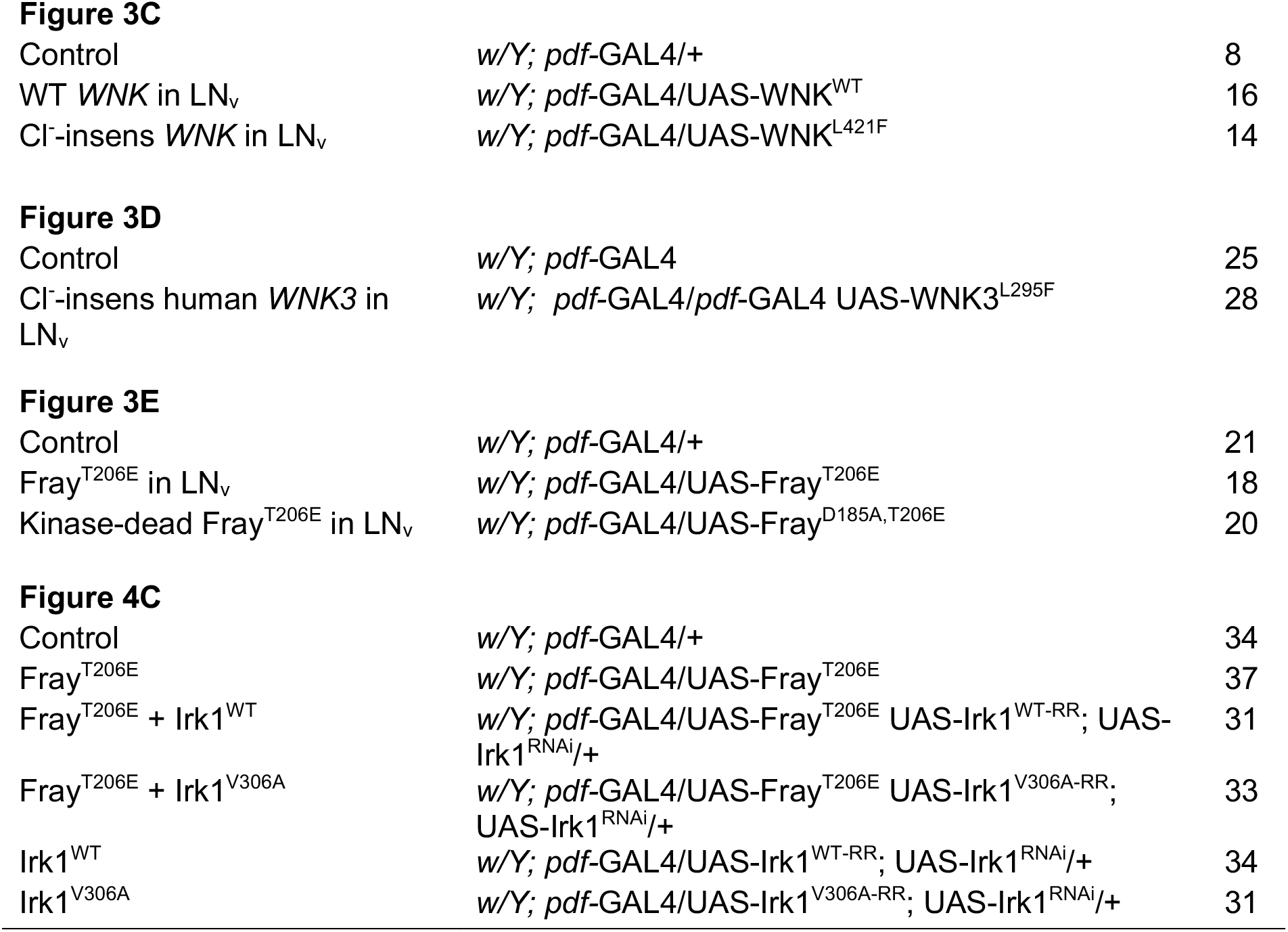
*Drosophila* genotypes. N=number of flies tested. For Figure 2A, n=number of cells analyzed.

| Name | Genotype | N |
| --- | --- | --- |
| **Figure 1A** |  |  |
| Control | *w/Y; pdf-GAL4 UAS-ClopHensor c202* | 14 |
| *Ncc69<sup>r2</sup>* | *w/Y; pdf-GAL4 UAS-ClopHensor c202; Ncc69<sup>r2</sup>* | 15 |
| **Figure 1B** |  |  |
| Control | *w/Y; Berlin* | 27 |
| *Ncc69<sup>r2</sup>/Ncc69<sup>r2</sup>* | *w/Y; Ncc69<sup>r2</sup>* | 22 |
| *Ncc69<sup>r2</sup>/Df* | *w/Y; Ncc69<sup>r2</sup>/ Df(3L)ED4475, p{3'.RS5+3.3}ED4475* | 24 |
| *Ncc69<sup>r2</sup>/+* | *w/Y; Ncc69<sup>r2</sup>/+* | 18 |
| *Df/+* | *w/Y; Df(3L)ED4475, p{3'.RS5+3.3}ED4475/+* | 30 |
| **Figure 1C** |  |  |
| Control | *w/Y; pdf-GAL4* | 29 |
| *Ncc69<sup>r2</sup>* | *w/Y; Ncc69<sup>r2</sup>* | 20 |
| LN<sub>v</sub> rescue | *w; pdf-GAL4; Ncc69<sup>r2</sup> UAS-HA-Ncc69<sup>2-3</sup>* | 23 |
| **Figure 1D** |  |  |
| Control | *w/Y; Berlin* | 15 |
| *Ncc69<sup>r2</sup>* | *w/Y; Ncc69<sup>r2</sup>* | 13 |
| **Figure 2A** |  |  |
| Control | *w/Y; Berlin* | 16 |
| *Ncc69<sup>r2</sup>* | *w/Y; Ncc69<sup>r2</sup>* | 13 |
| *Ncc69<sup>r2</sup> + het kcc mutant* | *w/Y; kcc<sup>DHS1</sup>/+; Ncc69<sup>r2</sup>* | 11 |
| **Figure 2B** |  |  |
| Control | *w/Y; pdf-GAL4* | 21 |
| *Ncc69<sup>r2</sup>* | *w/Y; Ncc69<sup>r2</sup>* | 12 |
| *Ncc69<sup>r2</sup> + LN<sub>v</sub> kcc knockdown* | *w/Y; pdf-GAL4/pdf-GAL4 UAS-kcc<sup>RNAi</sup>; Ncc69<sup>r2</sup>* | 16 |
| **Figure 3A** |  |  |
| Control | *w/Y; pdf-GAL4/+* | 15 |
| *Ncc69<sup>r2</sup>* | *w/Y; Ncc69<sup>r2</sup>* | 23 |
| *Ncc69<sup>r2</sup> + LN<sub>v</sub> WNK knockdown* | *w/Y; pdf-GAL4/UAS-WNK<sup>RNAi</sup>; Ncc69<sup>r2</sup> (UAS-WNK<sup>RNAi</sup> from Vienna, stock 106928)* | 28 |
| **Figure 3B** |  |  |
| Control | *w/Y; pdf-GAL4/+* | 9 |
| *Ncc69<sup>r2</sup>* | *w/Y; Ncc69<sup>r2</sup>* | 15 |
| *Ncc69<sup>r2</sup> + LN<sub>v</sub> Fray knockdown* | *w/Y; pdf-GAL4/+; UAS-fray<sup>RNAi</sup> Ncc69<sup>r2</sup>/Ncc69<sup>r2</sup> (UAS-fray<sup>RNAi</sup> from Rodan lab)* | 28 |
| **Figure 3C** |  |  |
| Control | *w/Y; pdf-GAL4/+* | 8 |
| WT *WNK* in LN<sub>v</sub> | *w/Y; pdf-GAL4/UAS-WNK<sup>WT</sup>* | 16 |
| Cl<sup>-</sup>-insens *WNK* in LN<sub>v</sub> | *w/Y; pdf-GAL4/UAS-WNK<sup>L421F</sup>* | 14 |
| **Figure 3D** |  |  |
| Control | *w/Y; pdf-GAL4* | 25 |
| Cl<sup>-</sup>-insens human *WNK3* in LN<sub>v</sub> | *w/Y; pdf-GAL4/pdf-GAL4 UAS-WNK3<sup>L295F</sup>* | 28 |
| **Figure 3E** |  |  |
| Control | *w/Y; pdf-GAL4/+* | 21 |
| Fray<sup>T206E</sup> in LN<sub>v</sub> | *w/Y; pdf-GAL4/UAS-Fray<sup>T206E</sup>* | 18 |
| Kinase-dead Fray<sup>T206E</sup> in LN<sub>v</sub> | *w/Y; pdf-GAL4/UAS-Fray<sup>D185A,T206E</sup>* | 20 |
| **Figure 4C** |  |  |
| Control | *w/Y; pdf-GAL4/+* | 34 |
| Fray<sup>T206E</sup> | *w/Y; pdf-GAL4/UAS-Fray<sup>T206E</sup>* | 37 |
| Fray<sup>T206E</sup> + Irk1<sup>WT</sup> | *w/Y; pdf-GAL4/UAS-Fray<sup>T206E</sup> UAS-Irk1<sup>WT-RR</sup>; UAS-Irk1<sup>RNAi/+</sup>* | 31 |
| Fray<sup>T206E</sup> + Irk1<sup>V306A</sup> | *w/Y; pdf-GAL4/UAS-Fray<sup>T206E</sup> UAS-Irk1<sup>V306A-RR</sup>; UAS-Irk1<sup>RNAi/+</sup>* | 33 |
| Irk1<sup>WT</sup> | *w/Y; pdf-GAL4/UAS-Irk1<sup>WT-RR</sup>; UAS-Irk1<sup>RNAi/+</sup>* | 34 |
| Irk1<sup>V306A</sup> | *w/Y; pdf-GAL4/UAS-Irk1<sup>V306A-RR</sup>; UAS-Irk1<sup>RNAi/+</sup>* | 31 |

**Supplemental Table 3, related to Figure 1.**
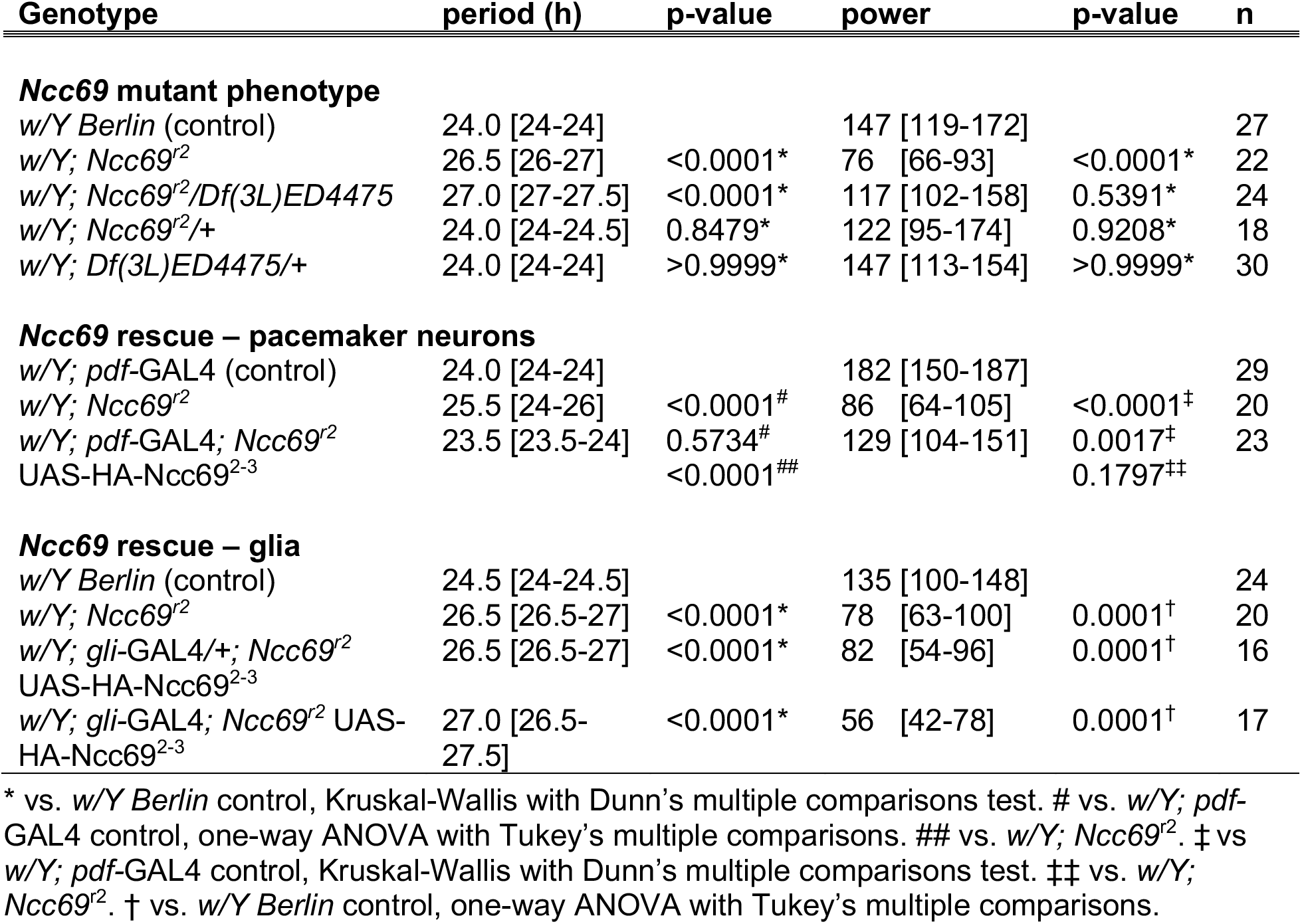
Period length and power of *Ncc69* mutants. All values shown are median [95% CI]. n=number of individual flies tested. *w;Ncc69^r2^* is a strong loss-of-function allele of the *Ncc69* NKCC. *Df(3L)ED4475* uncovers *Ncc69. Pdf-*GAL4 is expressed in the LNv pacemaker neurons. *Gli-*GAL4 is expressed in glia.

**Supplemental Table 4, related to Figure 3.**
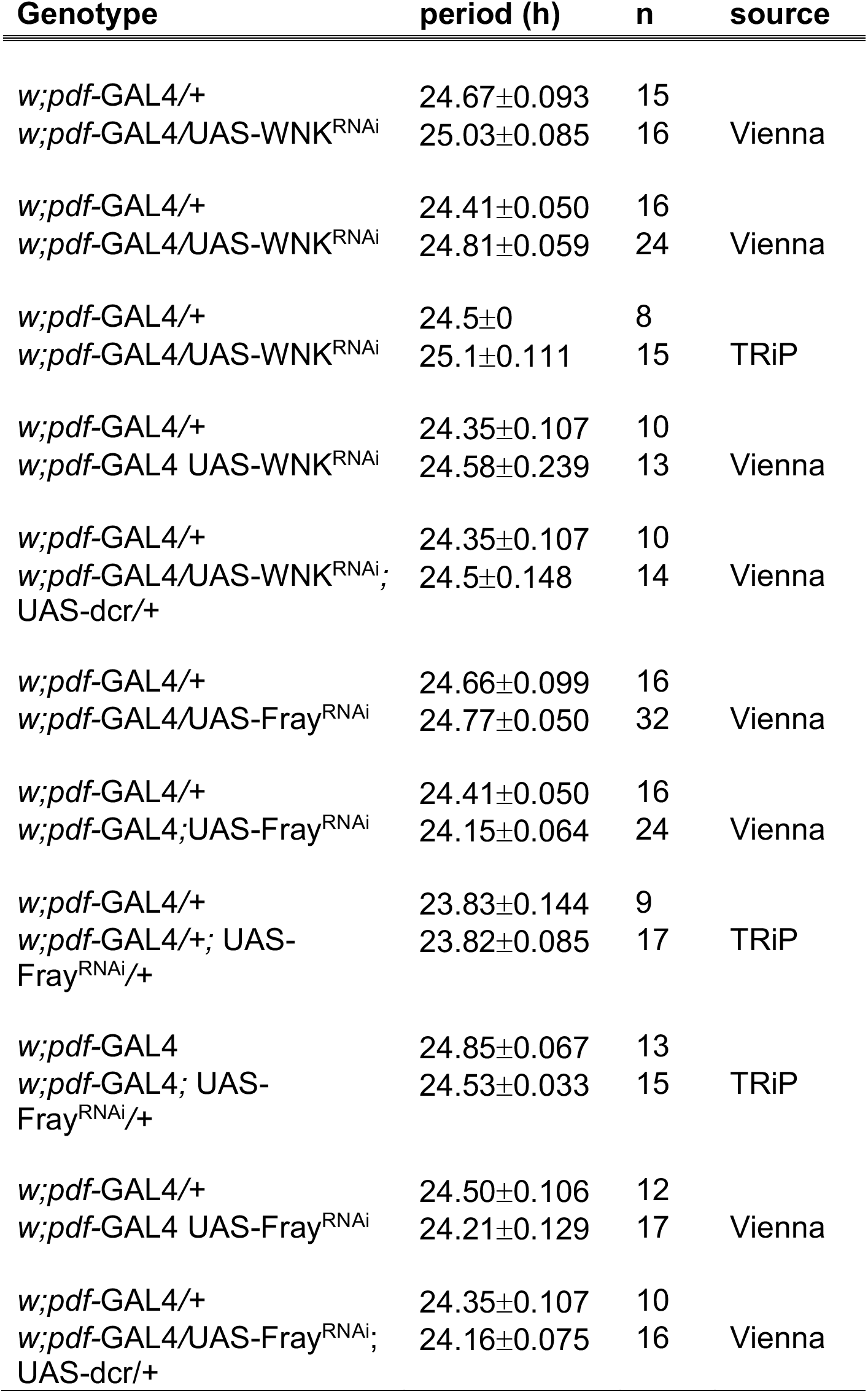
Effect of *WNK* and *Fray* knockdown on period length. All values shown are mean±SEM. n=number of individual flies tested. Each pair of genotypes is from an independent experiment. Source = source of RNAi line. *w;* UAS-WNK^RNAi^: Vienna = line 106928 from Vienna *Drosophila* Resource Center, TRiP = Transgenic Fly RNAi Project, Bloomington Stock Center #42521. *w;* UAS-Fray^RNAi^: Vienna = line 106919 from Vienna *Drosophila* Resource Center, TRiP = transgenic line made according to the methods of the Transgenic Fly RNAi Project, (Wu et al., 2014).

**Supplemental Figure 3, related to Figure 4.**
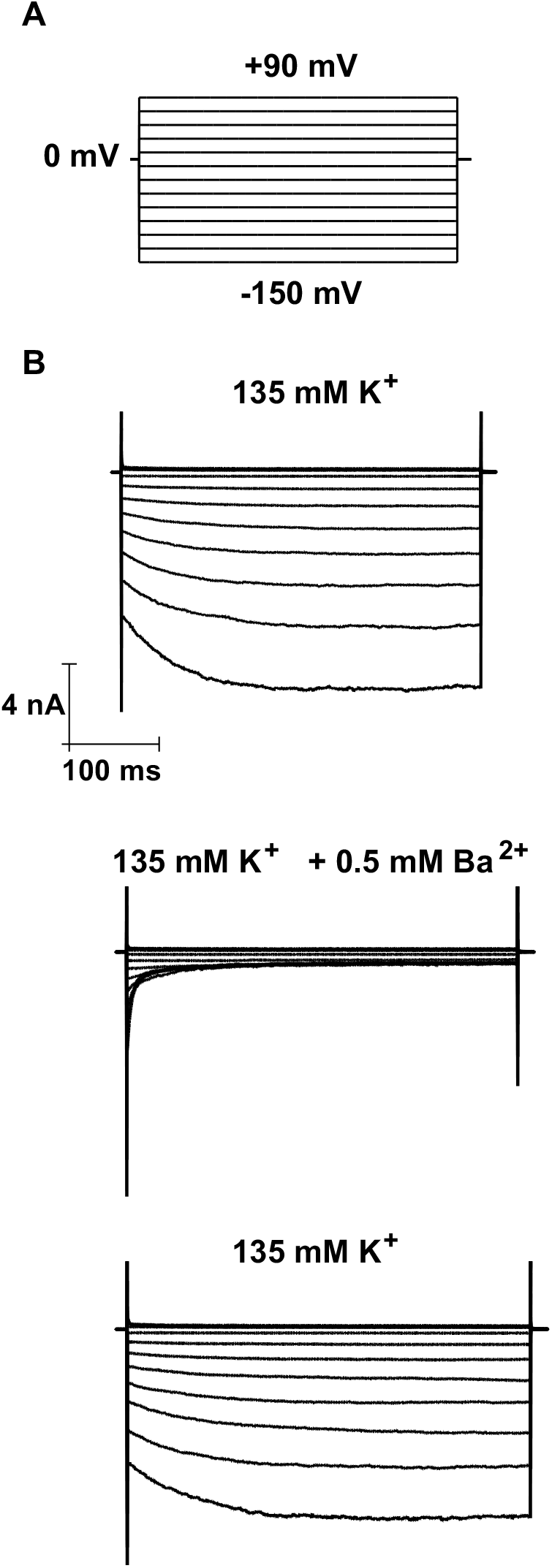
Measurement of barium-sensitive currents in Irk1-transfected S2-R+ *Drosophila* cultured cells. A) Stimulation protocol. Cells were held at 0 mV and stimulated for 400 ms with 20 mV step pulses from -150 mV to +90 mV. B) Traces showing real current in 135 mM K^+^, inhibited by 0.5 mM Ba^2+^ and recovered after washing in 135 mM K^+^. Ba^2+^-sensitive current density was compared in cells transfected with wild-type or mutant Irk1 with or without Fray^T206E^ co-expression (Figure 4).

**Supplemental Table 6.** *Drosophila melanogaster* strains.

| Genotype | Description | Source | Reference |
| --- | --- | --- | --- |
| *wBerlin* | Background/control strain | Rothenfluh |  |
| *w; pdf-GAL4* | LN<sub>v</sub> driver | Rosbash | (Renn et al., 1999) |
| *w; Ncc69<sup>r2</sup>* | *Ncc69* mutant | Leiserson | (Leiserson et al., 2010) |
| *w; Ncc69<sup>r2</sup> UAS-HA-Ncc69<sup>2-3</sup>* | *Ncc69* rescue | Leiserson | (Leiserson et al., 2010) |
| *w; gli-GAL4; Ncc69<sup>r2</sup> UAS-HA-Ncc69<sup>2-3</sup>* | Glial *Ncc69* rescue | Leiserson | (Leiserson et al., 2010) |
| *w; kcc<sup>DHS1</sup>* | *kcc* mutant | Hekmat-Scafe | (Hekmat-Scafe et al., 2006) |
| *w; UAS-kcc<sup>RNAi</sup>* | RNAi targeting *kcc* | VDRC 101742 | (Dietzl et al., 2007) |
| *w<sup>1118</sup>; Df(3L)ED4475, p{3'.RS5+3.3}ED4475/T M6C, cu<sup>1</sup> Sb<sup>1</sup>* | *Ncc69* deficiency | BDSC 8069 |  |
| *w; UAS-dcr-2* | Dicer transgene | BDSC 24651 | (Dietzl et al., 2007) |
| *w; UAS-WNK<sup>RNAi</sup>* | RNAi targeting *WNK* | BDSC 42521 | (Sun et al., 2018) |
| *w; UAS-WNK<sup>RNAi</sup> (Vienna)* | RNAi targeting *WNK* | VDRC 106928 | (Wu et al., 2014) |
| *w; UAS-WNK<sup>WT</sup>* | Wild-type *WNK* | Rodan | (Sun et al., 2018) |
| *w; UAS-WNK<sup>L421F</sup>* | Chloride-insensitive *WNK* | Rodan | (Sun et al., 2018) |
| *w; UAS-WNK3<sup>WT</sup>* | Human *WNK3* | Rodan | (Stenesen et al., 2019) |
| *w; UAS-WNK3<sup>L295F</sup>* | Chloride-insensitive human *WNK3* | Rodan | (Pleinis et al., 2021) |
| *w; UAS-fray<sup>RNAi</sup>* | RNAi targeting *Fray* | Rodan | (Wu et al., 2014) |
| *w; UAS-fray<sup>RNAi</sup> (Vienna)* | RNAi targeting *Fray* | VDRC 106919 | (Dietzl et al., 2007) |
| *w; UAS-Fray<sup>T206E</sup>* | Activated *Fray* | Rodan | (Wu et al., 2014) |
| *w; UAS-Fray<sup>D185A,T206E</sup>* | Kinase-dead activated *Fray* | This study |  |
| *w; UAS-Irk1<sup>RNAi</sup>* | RNAi targeting *Irk1* | BDSC 25823 | (Wu et al., 2015) |
| *w; UAS-Irk1<sup>WT-RR</sup>* | WT *Irk1* (RNAi-resistant) | This study |  |
| *w; UAS-Irk1<sup>V306A-RR</sup>* | V306A mutant *Irk1* (RNAi-resistant) | This study |  |
| *w; UAS-ClopHensor c202* | pH/chloride sensor | Krämer | (Sun et al., 2018) |

BDSC, Bloomington *Drosophila* Stock Center, Indiana University, Bloomington, IN
VDRC, Vienna *Drosophila* Resource Center, Vienna, Austria

**Supplemental Table 7.**
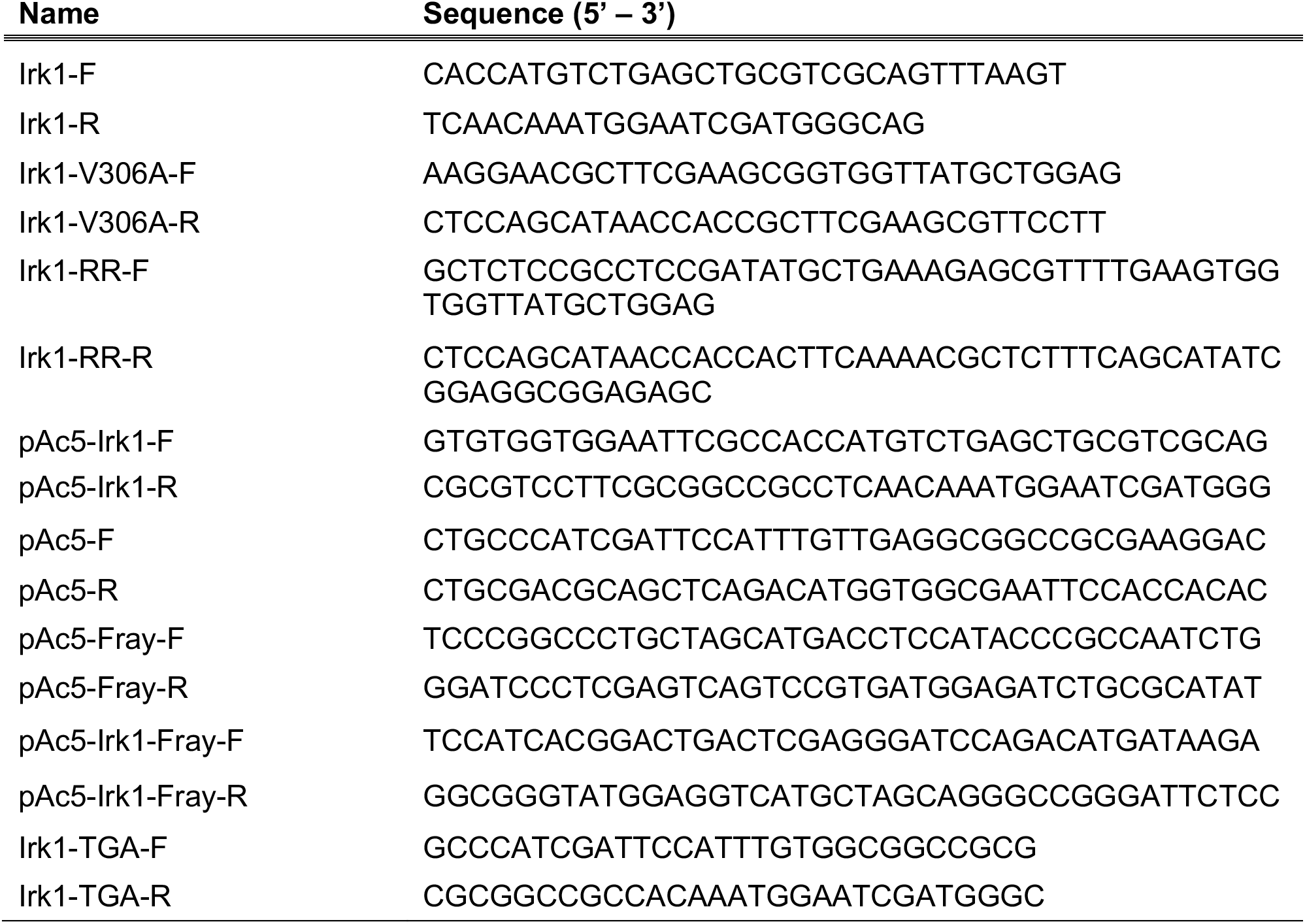
Primer sequences.

**Supplemental Table 8.** Newly generated plasmids.

| Name | Description |
| --- | --- |
| pENTR-Irk1<sup>WT</sup> | Irk1<sup>WT</sup> ORF in Gateway donor plasmid |
| pENTR-Irk1<sup>V306A</sup> | Irk1<sup>V306A</sup> ORF in Gateway donor plasmid |
| pENTR-Irk1<sup>WT-RR</sup> | RNAi-resistant Irk1<sup>WT</sup> ORF in Gateway donor plasmid |
| pENTR-Irk1<sup>V306A-RR</sup> | RNAi-resistant Irk1<sup>V306A</sup> ORF in Gateway donor plasmid |
| pUASg.attB-Irk1<sup>WT-RR</sup> | RNAi-resistant Irk1<sup>WT</sup> in pUAS plasmid |
| pUASg.attB-Irk1<sup>V306A-RR</sup> | RNAi-resistant Irk1<sup>V306A</sup> in pUAS plasmid |
| pIB-ClopHensor | pH/chloride sensor in S2 cell expression plasmid |
| pAc5-Irk1<sup>WT</sup> | Irk1<sup>WT</sup> in S2 cell expression plasmid |
| pAc5-Irk1<sup>V306A</sup> | Irk1<sup>V306A</sup> in S2 cell expression plasmid |
| pAc5-Irk1<sup>WT</sup>-Fray<sup>T206E</sup> | Irk1<sup>WT</sup> and Fray<sup>T206E</sup> in multi-cistronic S2 cell expression plasmid |
| pAc5-Irk1<sup>V306A</sup>-Fray<sup>T206E</sup> | Irk1<sup>V306A</sup> and Fray<sup>T206E</sup> in multi-cistronic S2 cell expression plasmid |

